# Axin2L regulates planar cell polarity in *Xenopus* embryos

**DOI:** 10.64898/2026.07.29.741488

**Authors:** Satheeja Santhi Velayudhan, Keiji Itoh, Sergei Y. Sokol

**Affiliations:** Department of Stem Cell Biology and Regenerative Medicine, Icahn School of Medicine at Mount Sinai, New York, NY, 10029 USA

**Author notes:** Correspondence: Sergei Y. Sokol.

**Keywords:** Axin2L, planar cell polarity, mechanosensation, neural tube closure, β-catenin, SSX2IP, ciliogenesis, wound healing

## Abstract

Axin proteins function as central scaffolds of the β-catenin destruction complex in the Wnt/β-catenin pathway that has been considered distinct from the planar cell polarity (PCP) pathway. Here, we show that dorsal expression of mutated Axin2L, a predominant Axin homolog in Xenopus early embryos, disrupts convergent extension movements and neural tube closure in a manner reminiscent of the mutations in vertebrate PCP genes. GFP-Axin2L becomes planar polarized in the epidermal cells adjacent to the folding neural plate and those surrounding a wound. Axin2L associated and colocalized with ADIP, a similarly polarized protein that is essential for neural tube morphogenesis and embryonic wound healing. Interference with Axin2L function abrogated Vangl2 accumulation at the anterior sides of neuroepithelial cells and impaired ciliogenesis in the gastrocoel roof plate. We propose that, in addition to the canonical Wnt pathway, Axin2L controls PCP signaling during vertebrate morphogenesis.

## Introduction

Wnt pathways play central roles in cell fate specification, cell proliferation and morphogenetic events in early embryonic development (MacDonald et al., 2009; Maurice and Angers, 2025; Nusse and Clevers, 2017). A critical regulatory step in the canonical Wnt/β-catenin pathway is mediated by the destruction complex of GSK3, APC and Axin that degrades β-catenin in the absence of a Wnt signal (Hart et al., 1998; Ikeda et al., 1998; Itoh et al., 1998). Wnt signals activate Dishevelled (Dvl) that interacts with Axin via the DIX (<u>Di</u>shevelled-A<u>x</u>in) domains to inactivate the destruction complex (Itoh et al., 2000; Kishida et al., 1999). This leads to the stabilization of β-catenin and transcriptional activation of Wnt target genes (MacDonald et al., 2009).

The upstream components of the canonical Wnt/β-catenin pathway including Frizzled (Fz) and Dishevelled (Dvl) also control planar cell polarity (PCP), coordinated orientation of epithelial cells in the plane of the tissue (Devenport, 2014; Goodrich and Strutt, 2011; McNeill, 2010; Peng and Axelrod, 2012; Zallen, 2007). Core PCP complexes consist of the Frizzled/Dvl/Diversin (Div or Ankrd6) and Vangl/Prickle (Pk) complexes, associated with Celsr/Flamingo. In vertebrates, PCP signaling is tightly connected to various morphogenetic movements, such as neural tube closure (Butler and Wallingford, 2017; Devenport, 2014; Williams and Solnica-Krezel, 2020). Downstream of Dvl, the β-catenin destruction complex is believed to be specific for the canonical Wnt pathway (Komiya and Habas, 2008; Semenov et al., 2007). Notably, the DIX domains of Dvl and Axin are distantly related to the domains found in plant polarity proteins (van Dop et al., 2020). Additionally, mutations in mouse Axin genes exhibit pleiotropic morphogenetic defects, including neural tube abnormalities (Aceves-Ewing et al., 2025; Qian et al., 2011; Yu et al., 2005; Zeng et al., 1997) that are characteristic for PCP signaling defects, raising a possibility that Axins may have broader roles in early development than previously anticipated.

This study evaluated a role of Axin2L, an abundant embryonic Axin homologue (Itoh et al., 2000) in PCP signaling using *Xenopus* embryos, an established experimental model for both canonical Wnt/β-catenin and PCP pathways (Hikasa and Sokol, 2013; Sokol, 2015). We find that Axin2L polarizes in the plane of the epidermal cells adjacent to the folding neural plate and the epithelium surrounding a wound. Axin2L interacts and colocalizes with Afadin DIL- and α-actinin-interacting protein (ADIP, also known as SSX2IP) (Asada et al., 2003) that is polarized in response to tension in a similar manner and is required for neural tube closure and wound healing (Chu et al., 2025). Interference with Axin2L function disrupted Vangl2 accumulation at the anterior sides of neuroepithelial cells, supporting the hypothesis that Axin2L regulates PCP in the vertebrate neural plate.

## RESULTS

### Axin2L-ΔRGS disrupts convergent extension movements in *Xenopus* embryos

Mouse and fish embryos with mutations in core PCP components commonly result in axial and neural tube abnormalities (Gray et al., 2011; Nikolopoulou et al., 2017; Torban and Sokol, 2021). Similarly, interference with PCP signaling in *Xenopus* embryos causes a characteristic phenotype resulting from disrupted convergent extension (CE) and neural tube defects (Sokol, 1996; Tada and Smith, 2000; Wallingford et al., 2000). Axin mutants also have neural tube and axial defects that were interpreted as the outcome of abnormal Wnt/β-catenin signaling. However, the strong phenotypes caused by deficient β-catenin signaling may obscure less prominent defects such as PCP abnormalities (Qian et al., 2011; Zeng et al., 1997). Our previous study suggested that Axin2L (formerly known as Xenopus Axin-related protein) may be involved in PCP signaling (Itoh et al., 2000) and the present study aimed at testing this hypothesis.

The deletion of the RGS domain in Axin1 and Axin2L was shown to have a dominant negative effect in different models, likely because it stabilizes β-catenin instead of degrading it (Fagotto et al., 1999; Itoh et al., 2000; Zeng et al., 1997). To characterize the effects of the Axin2L-ΔRGS construct on morphogenesis, four-cell embryos were injected dorsally with RNA encoding either full-length myc-Axin2L or myc-Axin2L-ΔRGS and scored at tailbud stages (**Fig. 1A, B**). Dorsal expression of full-length Axin2L caused strong ventralization, consistent with the known inhibitory activity of Axin proteins towards canonical Wnt/β-catenin signaling (**Fig. 1C, E**). These embryos failed to undergo normal axial development, supporting the idea that elevated Axin2L activity interferes with early dorsal patterning. By contrast, dorsal injection of Axin2L-ΔRGS RNA did not inhibit axial patterning but caused a dorsally-bent phenotype (**Fig. 1D, E**), characteristic of deficient CE movements (Sokol, 1996) and consistent with the possible involvement of Axin in PCP. Note that in our previous study, Axin2L-ΔRGS RNA has been targeted only to ventral blastomeres, in which it was shown to trigger secondary axis formation (Itoh et al., 2000).

**Fig. 1.**
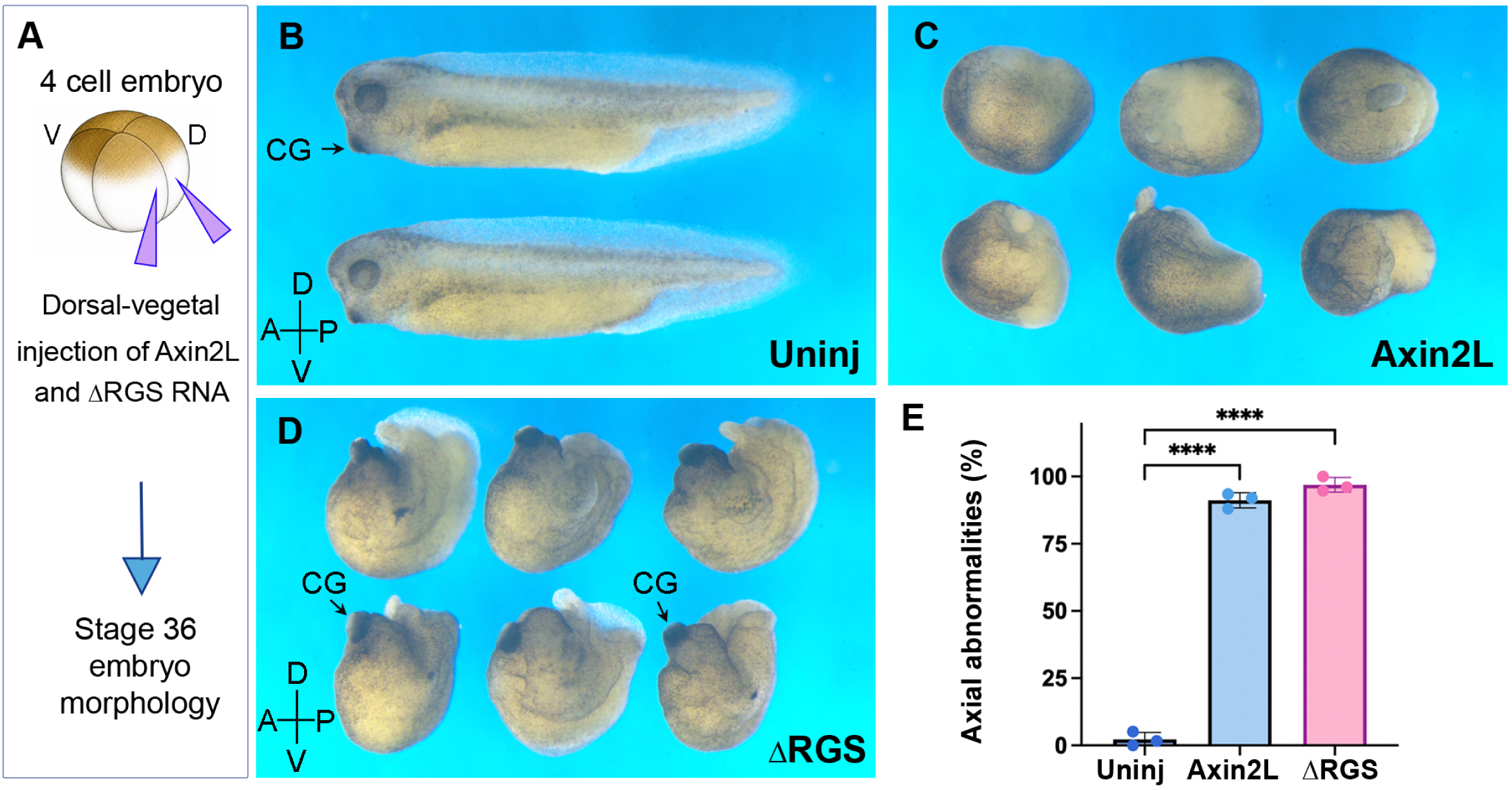
Axin2L with RGS domain deletion causes PCP-like defects. **(A)** Experimental scheme. Four-cell embryos were dorsally injected with 1 ng of myc-Axin2L or myc-Axin2L-ΔRGS RNA as indicated and their morphology was scored at stage 36. (**B**) Uninjected control embryos. (**C**) Ventralization of embryos expressing full-length myc-Axin2L. (**D**) Embryos expressing myc-Axin2L-ΔRGS exhibit shortened, dorsally bent phenotypes consistent with convergent extension defects. The anterior-posterior and dorsal-ventral axes are indicated. CG denotes the cement gland in (B, D). (**E**) Quantification of axial defects. Means ± s.d. are shown from three independent experiments, with 25-50 embryos per group. Welch’s t-test; ****p < 0.0001.

Together, these results indicate that full-length Axin2L and Axin2L-ΔRGS affect early *Xenopus* development in different ways. Axin2L strongly inhibits canonical Wnt-dependent axial patterning, whereas Axin2L-ΔRGS disrupts convergent extension movements during gastrulation and neural tube closure. These findings suggest that Axin2L participates in PCP-dependent morphogenesis in addition to its role in β-catenin signaling.

### Axin2L is polarized in the plane of the epidermis that is adjacent to the folding neural plate

Axin has been reported to distribute as cytoplasmic puncta or condensates that likely represent liquid-liquid phase separation (LLPS)(Fagotto et al., 1999; Itoh et al., 2000; Miete et al., 2022; Nong et al., 2021; Schaefer et al., 2018). Our recent discovery of the tension-dependent PCP axis that is perpendicular to the neural plate border (Chu et al., 2025) prompted us to test whether Axin2L condensates become planar polarized in this model. At stage 12, the puncta of GFP-Axin2L were broadly distributed within non-neural ectoderm cells without a detectable bias in tissue plane (**Fig. 2A-C**). After neural plate folding is initiated, we observed GFP-Axin2L accumulation in the proximal corners of cells that are adjacent to the neural plate border (**Fig. 2D-F)**. These cells are physically pulled by the neighboring apically constricting anterior neural ridge cells (Matsuda et al., 2023; Matsuda and Sokol, 2021), suggesting that Axin2L responds to mechanical forces during neural plate folding. This behavior resembles the planar polarization of ADIP and Diversin in the same tissue context (Chu et al., 2025; Velayudhan et al., 2025), supporting the idea that Axin2L is part of a mechanosensitive PCP protein complex that also includes ADIP and Diversin (see below). Consistent with this hypothesis, Diversin has been reported to bind Axin2 (Schwarz-Romond et al., 2002).

**Fig. 2.**
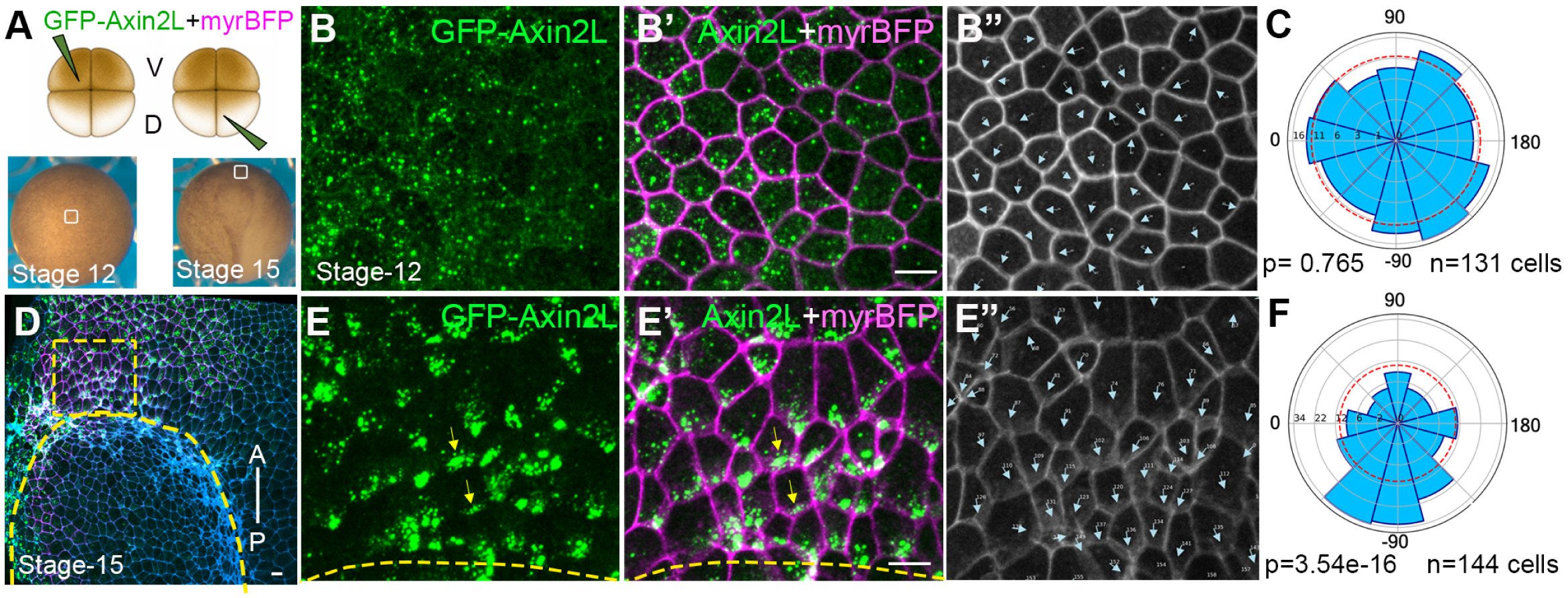
Planar polarization of Axin2L in ectoderm cells adjacent to the neural plate border. (**A**) Schematic of the experimental design. Embryos were injected with 400 pg GFP-Axin2L and 50 pg HA-myrBFP as a cell-boundary marker. White boxes indicate the regions imaged at stages 12 and 15. (**B–C**) At stage 12, GFP-Axin2L localizes as cytoplasmic puncta in gastrula ectodermal cells. (**B**) GFP-Axin2L signal alone. (**B′**) Merged image showing GFP-Axin2L with myrBFP-labeled cell boundaries. (**B″**) Cell segmentation and orientation-vector analysis showing random Axin2L puncta orientation. (**C**) Rose plot quantification showing no significant polarized orientation of Axin2L puncta in stage-12 ectodermal cells. p = 0.765, n = 131 cells. (**D**) Stage-15 Xenopus neurula embryo expressing GFP-Axin2L (green) and myrBFP (magenta), stained with phalloidin to mark F-actin (cyan). The neural plate border is outlined by the yellow dashed line, and the yellow dashed box indicates the region enlarged in panels E-E″. The anterior–posterior axis is indicated. (**E-F**) At stage 15, GFP-Axin2L becomes polarized in non-neural ectodermal cells adjacent to the folding neural tube. (**E**) GFP-Axin2L signal showing asymmetric enrichment of Axin2L puncta. Yellow arrows indicate polarized Axin2L accumulation. (**E′**) Merged image showing GFP-Axin2L with myrBFP-labeled cell boundaries. (**E″**) Cell segmentation and orientation-vector analysis showing biased Axin2L orientation toward the constricting neural ectodermal cells. (**F**) Rose plot showing significant Axin2L polarization at stage 15, with 90° corresponding to the anterior axis (*n* = 144 cells, P = 3.54 × 10⁻¹⁶). Chi-square test. Data are from three embryos per group in three independent experiments.

### Axin2L domain requirements for planar polarization during wound healing

The tension-dependent polarization of Axin2L was confirmed in a wound healing model. The actomyosin cable that forms around a wound repairs it via a ‘purse-string’ mechanism (Chu et al., 2025; Ossipova et al., 2014)(**Fig. 3A, B**). Before wounding, GFP-Axin2L condensates were distributed within the cytoplasm in a non-polarized manner (**Fig. 3C, E).** Manual wounding of the epidermis in embryos injected with GFP-Axin2L resulted in the accumulation of Axin2L at the cell edges that are proximal to the wound (**Fig. 3D, F).** This result confirms the tension-dependent planar polarization of Axin2L.

**Fig. 3.**
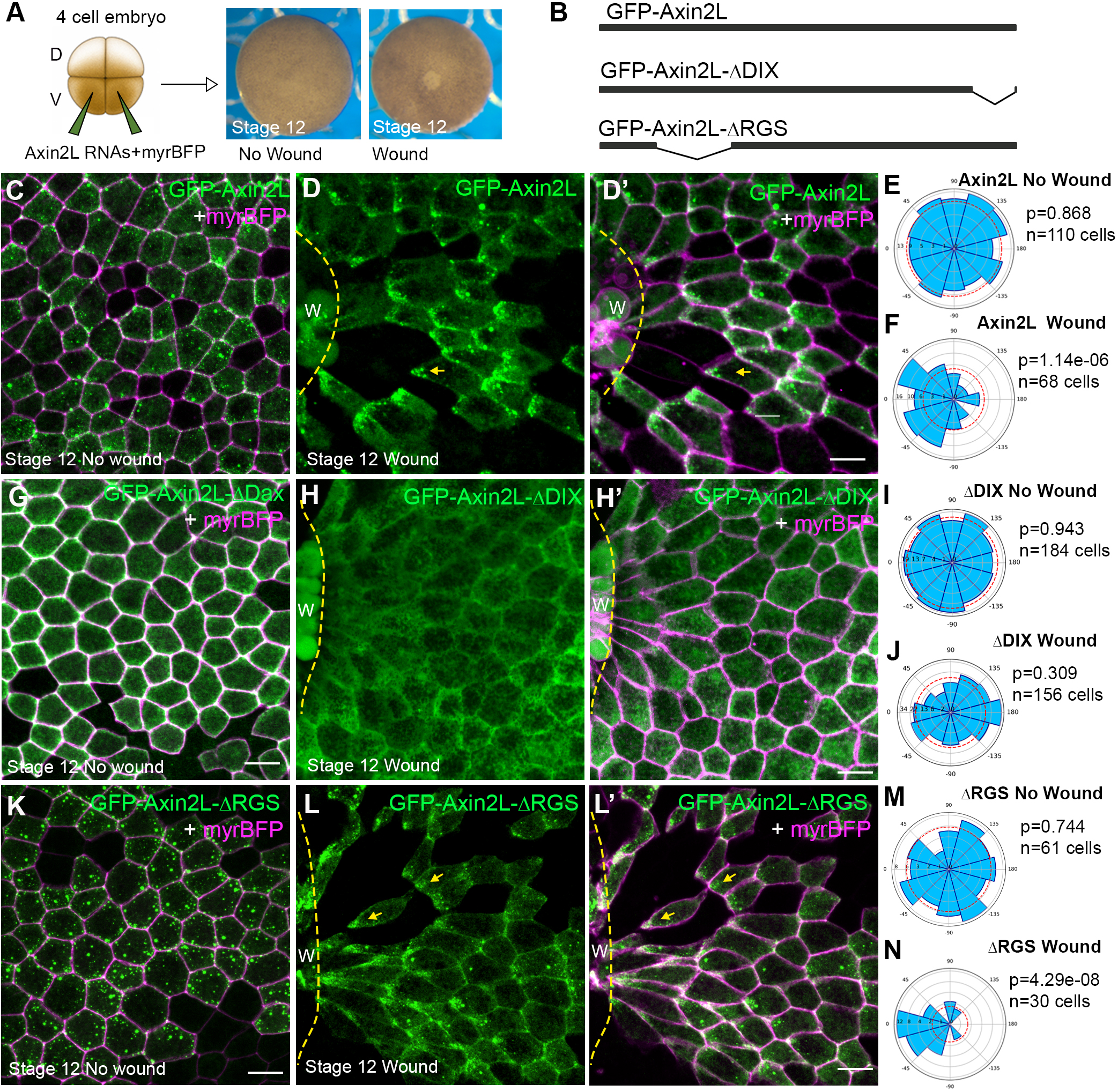
Polarization of Axin2L constructs in plane of the epithelium surrounding a wound. **(A)** Experimental scheme. Four-cell stage embryos were injected with RNA encoding GFP-Axin2L, GFP-Axin2L-ΔDIX, or GFP-Axin2L-ΔRGS (1.5 ng) together with myrBFP RNA (50 pg) as a cell border tracer. Stage 12 (st.12) surface ectoderm was imaged intact or at 30-45 min after wounding. **(B)** Schematic of GFP-Axin2L constructs used in this experiment. GFP-Axin2L-ΔDIX lacks the C-terminal DIX domain, whereas GFP-Axin2L-ΔRGS lacks the RGS domain. **(C)** Localization of GFP-Axin2L puncta in control st. 12 ectoderm. **(D, D′)** After wounding, GFP-Axin2L puncta become asymmetrically enriched in the cell corners proximal to the wound edge. **(E, F)** Rose plots reflect data in (C, D). **(G)** Diffuse cytoplasmic and cortical localization of GFP-Axin2L-ΔDIX in control st.12 ectoderm. **(H, H′)** No obvious enrichment of GFP-Axin2L-ΔDIX toward wound edge**. (I, J)** Rose plots are related to data in (G, H). **(K)** Puncta of GFP-Axin2L-ΔRGS in control ectoderm. **(L, L′)** After wounding, GFP-Axin2L-ΔRGS puncta polarize toward wound edge. **(M, N)** Rose plots quantify data in (K, L). GFP-Axin2L proteins are in green, myrBFP is in magenta. Yellow dashed lines indicate wound edge, yellow arrows indicate polarized protein accumulation. Size bars, 20 µm. Rose plots quantify Axin2L puncta orientation relative to the animal pole in control ectoderm or wound edge in wounded ectoderm, with 0° indicating the reference direction. Chi-square test. Data are from three embryos per group, three independent experiments.

This assay has been used to identify the domains of Axin2L required for its polarization. Several Axin2L constructs with various domain deletions have been made **(Fig. S1A)** and found to be expressed at comparable levels **(Fig. S1B-D)**. Deletion of the DIX domain reduced Axin2L puncta and produced a diffuse cytoplasmic distribution both before or after wounding (**Fig. 3G–J**). This mutant failed to polarize toward the wound margin (**Fig. 3J**), indicating that the DIX domain is required for both Axin2L condensate formation and wound-induced planar polarization. By contrast, the deletion of the RGS domain did not prevent wound-directed polarization. GFP-Axin2L-ΔRGS was randomly distributed in unwounded ectoderm (**Fig. 3K, M**) but significantly polarized toward the wound margin after wounding (**Fig. 3L,L′, N**). Additional deletions showed that the Tankyrase- and β-catenin-binding domain were also dispensable for wound-induced Axin2L polarization (**Fig. S2A-E, J-M**). By contrast, deletion of the GSK3β-binding domain (GBD) did not abolish puncta formation but prevented wound-directed polarization (**Fig. S2F–I**).

Together, these results show that the DIX domain and the GBD are required for wound-induced directional polarization but only the DIX domain is necessary for condensate formation. In contrast, GFP-Axin2L constructs lacking the RGS, β-catenin-binding, and Tankyrase-binding domains still polarized in our model similarly to full-length Axin2L.

### Axin2L associates and colocalizes with ADIP in embryonic ectoderm

ADIP is a mechanosensitive microtubule-associated protein that is essential for neural tube morphogenesis and embryonic wound healing (Chu et al., 2025). Since Axin2L localization in the tissue under tension was similar to that of ADIP, we tested whether these proteins might be present in the same mechanosensitive complexes. GFP-Axin2L was expressed in the anterior neural ectoderm (**Fig. 4A**) and showed polarized puncta distribution (**Fig. 4B, D**). This distribution matched the planar-polarized localization of HA-RFP-ADIP in the same tissue region (**Fig. 4C, E**). Coexpression of GFP-Axin2L and HA-RFP-ADIP revealed overlapping puncta and planar polarized distribution of both proteins in the anterior neural ectoderm (**Fig. 4F-H, Fig. S3**). Of note, 75% of Axin2L puncta contained ADIP (**Fig. S3**).

**Fig. 4.**
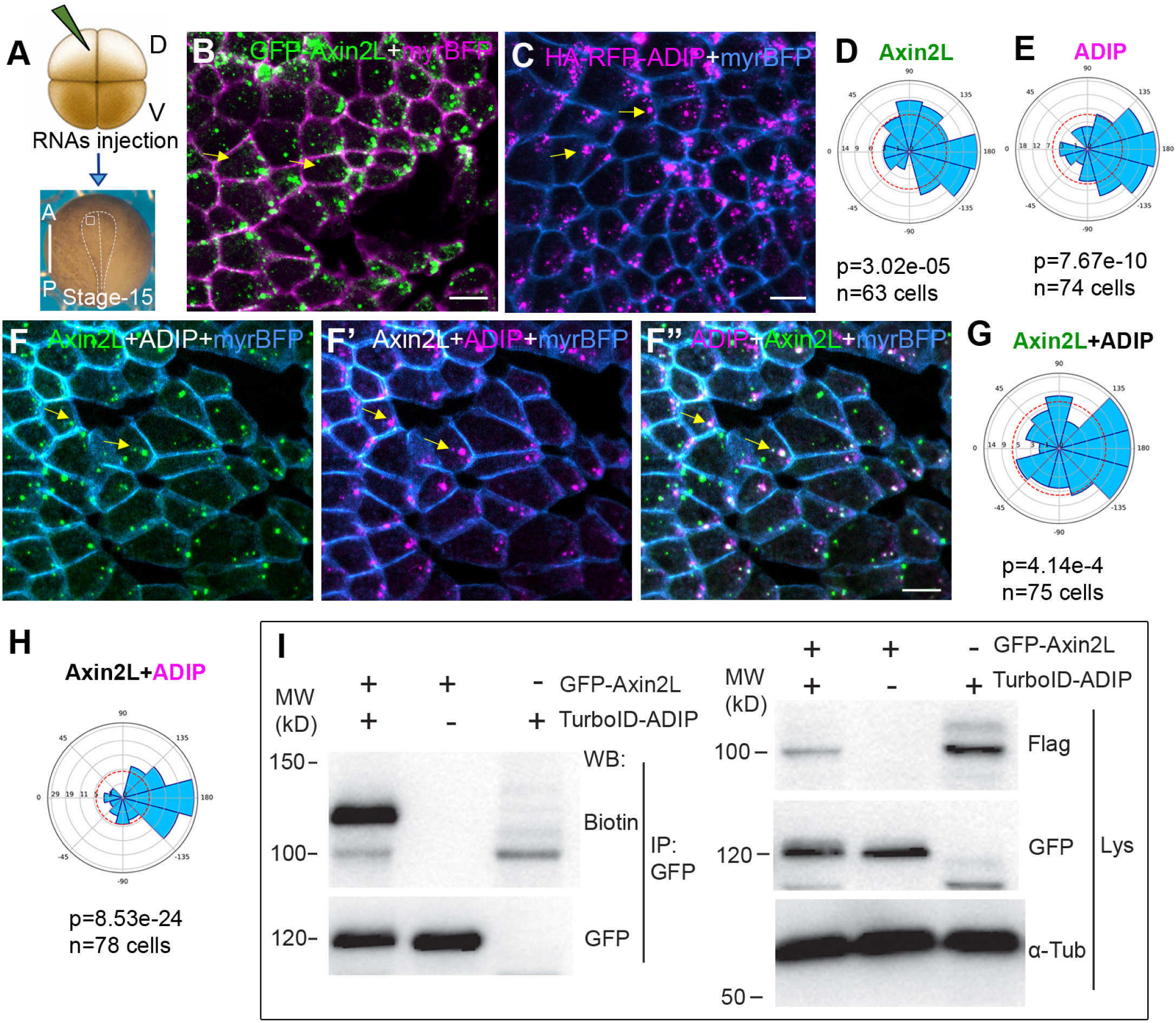
Axin2L and ADIP puncta colocalize and planar polarize toward neural plate midline. (**A)** Injection scheme. Four-cell embryos were injected dorsally with GFP-Axin2L (300 pg), HA-RFP-ADIP (300 pg) and myrBFP RNA (50 pg) as indicated and imaged at stage 15. GFP-Axin2L is in green, HA-RFP-ADIP in magenta, and myrBFP in cyan or magenta, as indicated. Scale bars, 20 μm. (**B)** Polarized GFP-Axin2L puncta (yellow arrows) in anterior neural ectoderm with myrBFP marking cell borders. (**C**) Polarization of HA-RFP-ADIP puncta in anterior neural ectoderm (yellow arrows). (**D, E)** Rose plots quantify individually expressed GFP-Axin2L (**D**) and HA-RFP-ADIP (**E**) puncta orientation. (**F–F″**) Coexpression of GFP-Axin2L and HA-RFP-ADIP. (**F**) polarized GFP-Axin2L puncta with myrBFP-marked cell borders. **F′**, Polarized HA-RFP-ADIP puncta with myrBFP-marked cell borders. **F″**, Merged image with overlapping Axin2L and ADIP puncta (yellow arrows). (**G, H**) Rose plots depict GFP-Axin2L (**G**) and HA-RFP-ADIP (**H**) puncta orientation in cells coexpressing Axin2L and ADIP. Rose plots in (**D, E, G, H**) show polarity angles, with 90° corresponding to the anterior axis of the embryo. Numbers of analyzed cells and p values are indicated in each plot. Chi-square test, data are from three embryos per group, three experiments. (**I**) Proximity biotinylation of GFP-Axin2L by TurboID-ADIP. Embryos were injected with GFP-Axin2LΔTBD (400 pg) and Flag-TurboID-ADIP (200 pg) RNAs, and 0.8 mM Biotin. Biotinylation was detected in GFP-Axin2L after pulldowns with GFP trap (left panel). Expression levels of Flag-TurboID-ADIP and GFP-Axin2L in lysates are shown (right panel). αActin serves as loading control.

In traditional coprecipitation assays, ADIP was not effectively pulled down with Axin2L, possibly because the complex was transient or sensitive to the presence of detergents. We therefore tested for the proximity of the two proteins using an ADIP-TurboID biotinylation assay (Branon et al., 2018). When GFP-Axin2L was pulled down from embryos expressing TurboID-ADIP, its biotinylation was easily detected by immunoblotting (**Fig. 4I**). This result indicates that Axin2L is in close proximity to ADIP in embryonic cells. Together our observations suggest that ADIP and Axin2L can be present in the same mechanosensitive protein complex.

### Axin2L is required for anterior polarization of Vangl2 in neuroepithelial cells

We next asked whether embryonic PCP in the neural plate is affected by Axin2L-ΔRGS construct. We evaluated endogenous Vangl2 that is accumulated at anterior cell edges in the *Xenopus* neural plate and can be easily detected with home-made antibodies (Ossipova et al., 2015). Full-length Axin2L did not significantly affect Vangl2 anterior accumulation, whereas the Axin2L-ΔRGS construct had a strong inhibitory effect **(Fig. 5A-E).** Similar effect was observed using a different Axin construct, Axin2L-C (**Fig. S4**). Thus, interference with Axin2L function leads to the disruption of endogenous Vangl2 planar polarity.

**Fig. 5.**
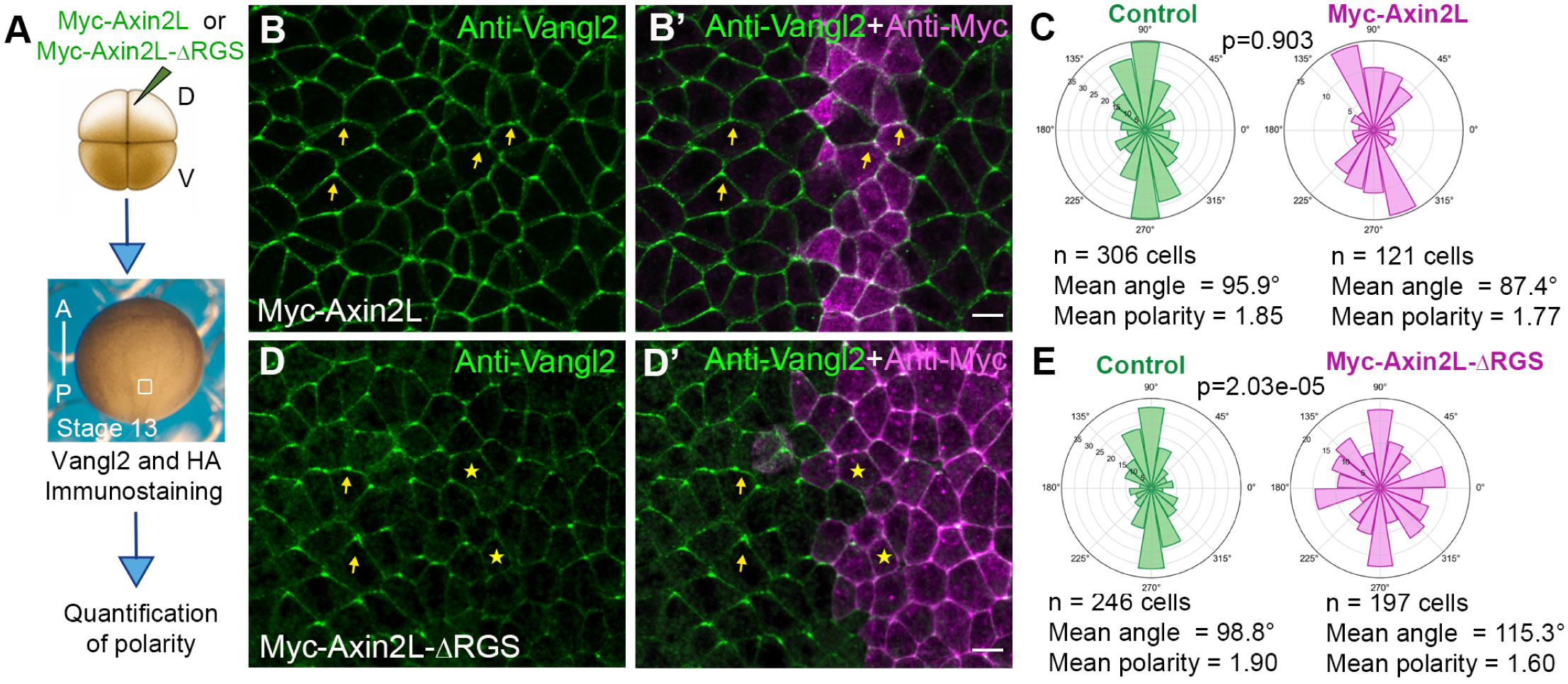
Interference with Axin2L function interferes with endogenous Vangl2 polarization in the neural plate. (**A**) Experimental scheme. Four-cell embryos were injected into one dorsal animal blastomere with 500 pg RNAs encoding myc-Axin2L or myc-Axin2L-ΔRGS. Embryos were fixed at stage 13 and immunostained for endogenous Vangl2 (green) and Myc epitopes (magenta) to identify injected cells. Anterior-posterior (A-P) axis is indicated. (**B, B′**) Embryos expressing myc-Axin2L show anteriorly enriched Vangl2 similar to neighboring uninjected cells. (**C**) Rose plots quantify data shown in B, B’ lacking significant effect on Vangl2 distribution as compared to control cells. (**D, D′)** Embryos expressing myc-Axin2L-ΔRGS show reduced or disrupted anterior Vangl2 enrichment in injected cells. Yellow arrows indicate anteriorly biased Vangl2 localization in control cells, whereas yellow asterisks mark cells with disrupted Vangl2 polarity. (B, D) Scale bars, 20 μm. (**E**) Rose plots quantification data shown in D, D’ with altered Vangl2 distribution in myc-Axin2L-ΔRGS-expressing cells. (C, E) Distribution of Vangl2 polarity axes determined by quadrant-ratio analysis, with 90° corresponding to the anterior direction. The number of analyzed cells, mean polarity angles and p values, are indicated (two-sided Mann-Whitney test). Three embryos per group, three experiments.

We also assessed Vangl2 polarity in cells depleted of Axin2L using specifically designed morpholinos (MOs). MO efficacy and specificity was validated with home-made antibodies to Axin2L (Alexandrova and Sokol, 2010)(**Fig. S5**). Control MO-injected cells retained anteriorly biased endogenous Vangl2 localization in the neural plate (**Fig. 6A-C**). By contrast, Axin2L-MO-injected cells showed reduction of anterior Vangl2 accumulation and significant disruption of the polarity axis (**Fig. 6D-E**). This defect was partly but consistently rescued by GFP-Axin2L-ΔTBD RNA that lacks MO target sequence (**Fig. 6F-G**). Together, these results indicate that Axin2L is required for the anterior accumulation of endogenous Vangl2 in neuroepithelial cells. This finding supports a functional role for Axin2L in PCP-dependent neural plate morphogenesis.

**Fig. 6.**
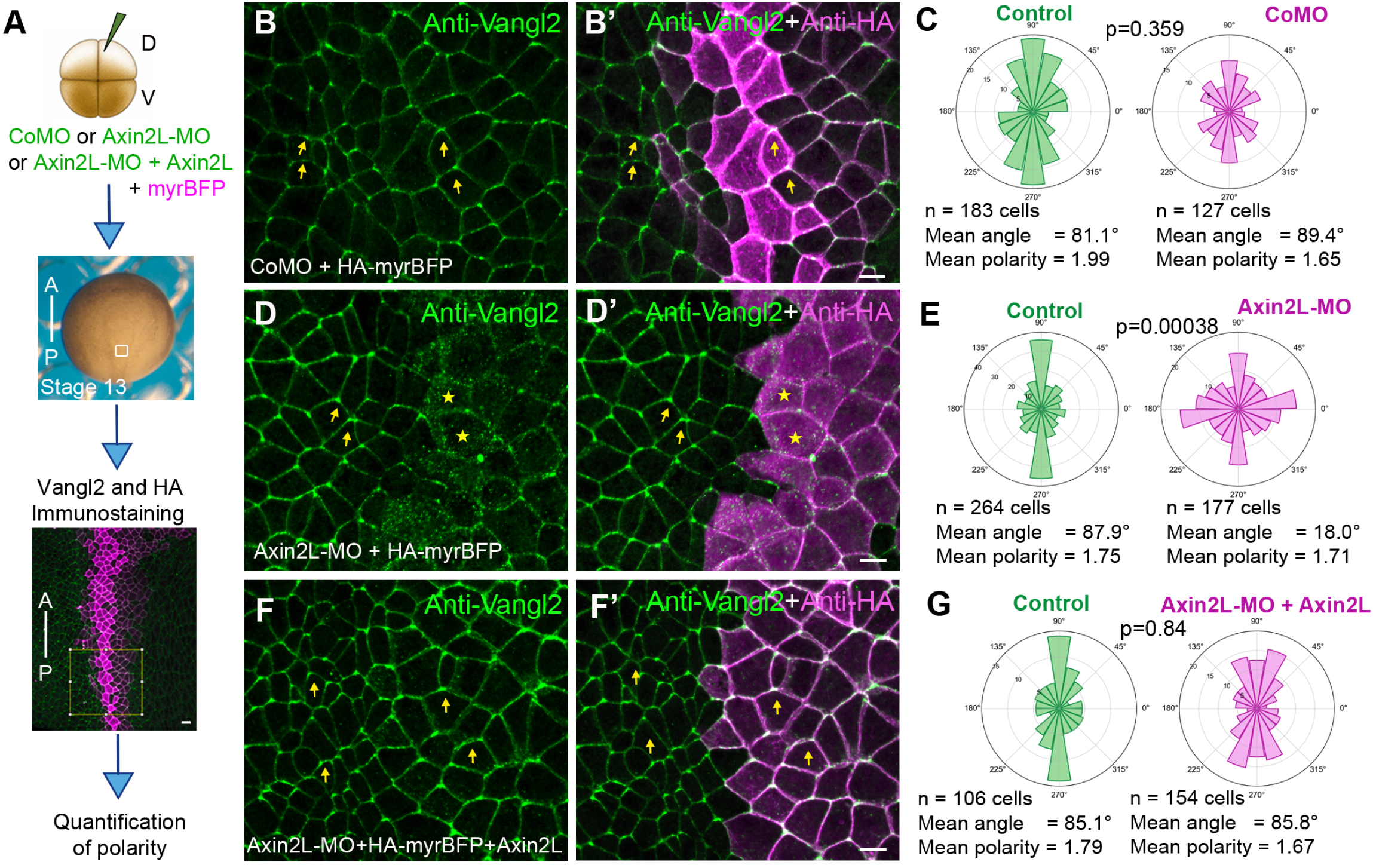
Axin2L depletion disrupts Vangl2 planar polarity. (**A**) Experimental scheme. Four-cell embryos were injected into one dorsal animal blastomere with control morpholino (CoMO, 30 ng), Axin2L MO (30 ng), or Axin2L-MO (30 ng) together with 500 pg Axin2L RNA and 50 pg HA-myrBFP as a lineage tracer. Fixed embryos, stage 13, were immunostained for endogenous Vangl2 (green) and HA epitopes (magenta) to identify injected cells. Anterior-posterior (A-P) axis is indicated. (**B, B**′) CoMO-injected cells retain anteriorly enriched Vangl2, similar to neighboring uninjected control cells. (**C**) Quantification of data in B, B′. (**D, D**′) Axin2L-MO-injected cells have disrupted anterior Vangl2 enrichment compared with neighboring control cells. Yellow arrows indicate anteriorly biased Vangl2 localization in control cells, whereas yellow asterisks mark cells with disrupted Vangl2 polarity. (B, D, F) Scale bars, 20 μm. (**E**) Quantification of data shown in D, D′. (**F, F**′) Coexpression of Axin2L-ΔTBD RNA with Axin2L-MO restores anteriorly enriched Vangl2 localization in injected cells. (**G**) Quantification of data shown in F, F′. (C, E, G) Distribution of Vangl2 polarity axes determined by quadrant-ratio analysis, with 90° corresponding to the anterior direction. Number of analyzed cells, mean polarity angles and p values are indicated. AP/ML polarity ratios were compared using a two-sided Mann–Whitney test. Three embryos per group, three experiments.

### Axin2L is required for ciliogenesis in the gastrocoel roof plate

Many Wnt and PCP signaling components are necessary for cilia formation and function (Adler and Wallingford, 2017; Niehrs et al., 2025; Park et al., 2006; Wallingford and Mitchell, 2011). We thus assessed whether Axin2L plays a role in ciliogenesis in the *Xenopus* gastrocoel roof plate (GRP), a tissue equivalent to the ciliated zebrafish Kupffer vesicle and mouse posterior node (Blum et al., 2014). Embryos were injected dorsally with Axin2L-MO and monocilia in the GRP were visualized by acetylated tubulin staining (**Fig. 7A, B**). Control cells displayed long cilia in the GRP. In contrast, interference with Axin2L resulted in absent or short cilia (**Fig. 7C, D**). Together, these findings indicate that Axin2L is required for monocilia elongation in the GRP. Combined with the disruption of endogenous Vangl2 polarity in Axin2L-deficient neural plate cells, these findings support a role for Axin2L in PCP-associated morphogenesis and ciliogenesis in *Xenopus* early embryos.

**Fig. 7.**
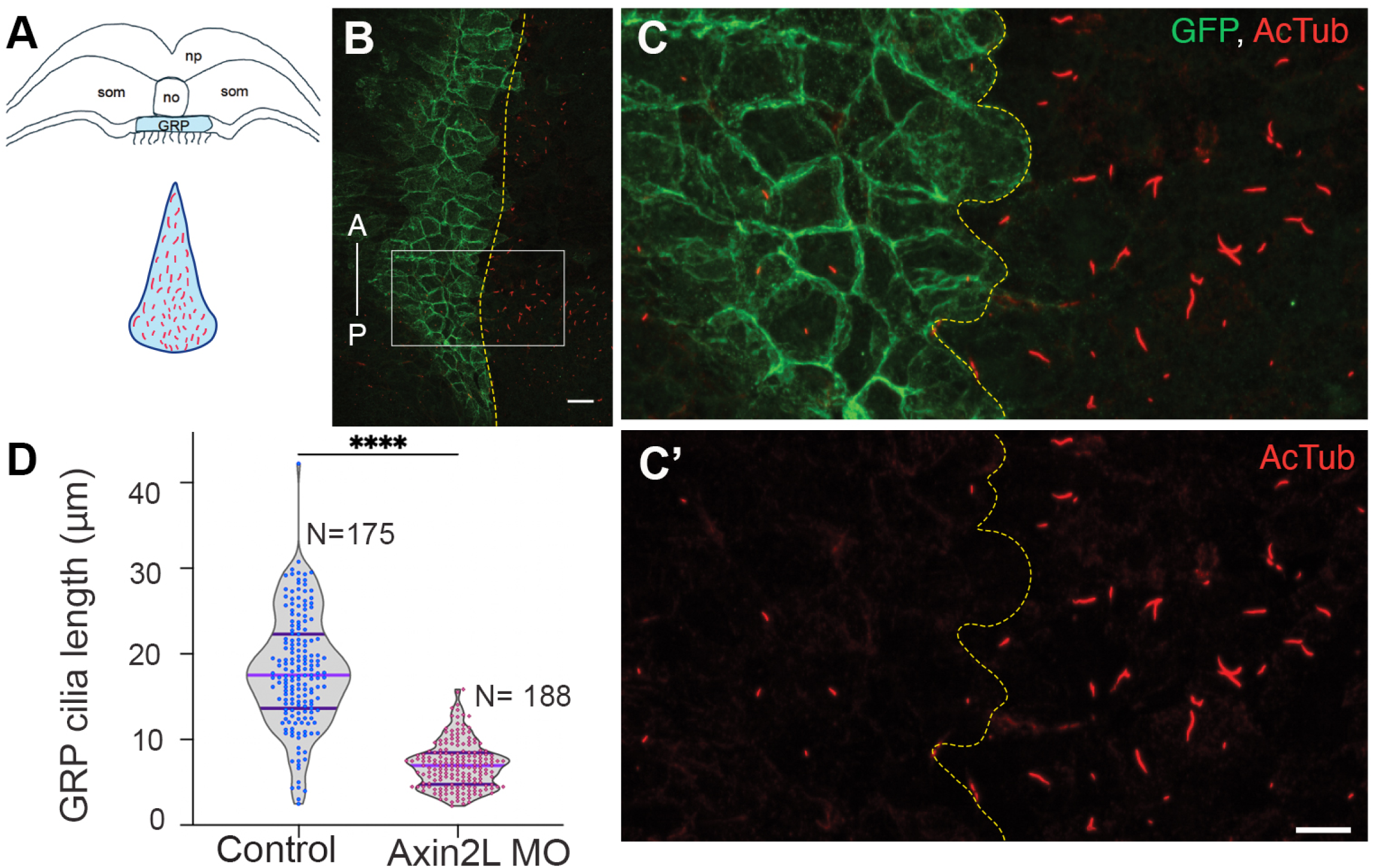
Disruption of the Axin2L function inhibits cilia formation in the gastrocoel roof plate. Axin2L MO (40 ng) and myrGFP RNA (10 pg) were coinjected into one dorsal blastomere at the medial equator of four-cell embryos. Gastrocoel roof plate (GRP) was dissected at stage 17 and immunostained with anti-acetylated Tubulin (AcTub) and anti-GFP antibodies. (**A**) GRP scheme with cross-section (top) and *en face* views. (**B**) Immunostained GRP. Anterior (A) to posterior (P) axis is indicated. Midline is indicated with dashed line. Box in B was magnified in (**C, C’**) Scale bars in B and C, C’ is 200 µm and 50 µm respectively. Ventral midline is indicated by dashed lines. (**D**) Quantification of cilia length in GFP-positive Axin2L-MO-injected and GFP-negative control cells. Cilia length was scored in eight GRP explants. N, numbers of scored cilia. Student’s t test, **** p <0.0001. Violin plots indicate medians and quartiles.

## DISCUSSION

After the original discovery of Axin protein in 1997, many studies culminated in the model, in which Axin is an important scaffold in the β-catenin degradation complex, the central regulatory point in canonical Wnt signaling (Behrens et al., 1998; Hart et al., 1998; Ikeda et al., 1998; Itoh et al., 2000; Itoh et al., 1998; Nusse and Clevers, 2017). Our results warrant a revision of the Axin function in early development. We find that Axin2L becomes polarized in the plane of the *Xenopus* embryonic epidermis near the neural plate border and adjacent to a wound. Besides exhibiting PCP-like distribution to cell edges proximal to the source of unidirectional pulling forces, two Axin2L truncated constructs inhibited convergent extension movements in the embryo, a phenotype characteristic of PCP defects. Interference with the Axin2L function disrupted the normal accumulation of Vangl2 at anterior cell corners in the neural plate. Based on our observations, we propose that Axin2L is a component of tension-sensitive PCP, which we recently characterized for ADIP, another scaffolding protein (Asada et al., 2003; Chu et al., 2025; Velayudhan et al., 2025).

Our findings do not support an exclusive role of the destruction complex in canonical Wnt signaling and imply that Axin2L also participates in tension-sensitive PCP. This function agrees well with previously found interactions of Axin with Dvl, a core PCP component (Itoh et al., 2000; Kishida et al., 1999; Smalley et al., 1999), and YAP, Hippo pathway mediator (Azzolin et al., 2014). Notably, consistent with their role in cell polarity, both Axin and Dvl function to polarize growing dendrites (Weiner et al., 2020) and reorient the centrosome and microtubules in the wound scratch assay (Schlessinger et al., 2007), supporting the proposed role for PCP signaling in wound healing (Caddy et al., 2010; Chu et al., 2025).

Our findings broaden the developmental role of Axin2L beyond canonical Wnt/β-catenin regulation and suggest that Axin2L participates in a mechanosensitive PCP network linking cortical polarity, tissue morphogenesis, and ciliogenesis. Similarly, the Wnt coreceptor LRP5/6 was proposed to function in both canonical Wnt/β-catenin and PCP signaling pathway (Weber et al., 2025). Mechanistically, the analysis of Axin2L domains points to the requirement of the Axin DIX domain and the GBD for protein polarization. Of note, distant homologues of the DIX domains in *Arabidopsis* are involved in plant tissue polarity (van Dop et al., 2020). The significance of the GBD for Axin2L polarization is currently unclear but may relate to its ability to bind Diversin (Schwarz-Romond et al., 2002). Whereas the RGS domain is not required for polarization of Axin2L, it is likely required for its function in PCP signaling, possibly via its interaction with APC, β-catenin or Gα proteins (Castellone et al., 2005; Itoh et al., 1998). The cortical asymmetry of β-catenin in *C. elegans* (Takeshita and Sawa, 2005) supports the view that Axin2L could play a role in local β-catenin degradation on one side of the cell. Future studies are needed to test the functional significance of β-catenin degradation at the cortex during Axin2L polarized accumulation, as compared to the well known transcriptional activity of β-catenin in the nucleus.

All proteins composing the observed tension-sensitive PCP complex, including Axin, Div, ADIP and Dvl, are known to associate with the centrosome (Alexandrova and Sokol, 2010; Cervenka et al., 2016; Fumoto et al., 2009; Hori et al., 2015; Itoh et al., 2009), indicating their potential role in centrosome-related functions, such as ciliogenesis. Indeed, we show that Axin2L is required for cilia growth in the GRP, supporting previous evidence reported for Dvl, Div and ADIP (Hashimoto et al., 2010; Hori et al., 2014; Park et al., 2008; Shnitsar et al., 2015; Yasunaga et al., 2011). The association of Axin2L with ADIP observed in this study and the previously reported binding of Axin2 to Dvl and Diversin (Fagotto et al., 1999; Itoh et al., 2000; Kishida et al., 1999; Schwarz-Romond et al., 2002; Smalley et al., 1999) suggest that the molecules are polarized using the same mechanism. One possibility is that these proteins coalesce into and function as LLPS condensates that are nucleated by the ADIP-containing pericentriolar material. Indeed, both Axin and Dvl puncta were reported to be LLPS condensates (Kang et al., 2022; Nong et al., 2021; Schubert et al., 2022; Vamadevan et al., 2022), and the centrosome was proposed to nucleate the β-catenin destruction complex (Itoh et al., 2009; Lach et al., 2022). Additional work is needed to investigate underlying mechanisms and define how the movement of these PCP condensates in the cell is regulated by tension.

## Materials and Methods

### Plasmid constructs, RNAs and morpholinos

The plasmids used in this study were pXT7-Myc-Axin2L, pXT7-Myc-ΔRGS-Axin2L, pXT7-HA-Axin2L, pXT7-HA-Axin2L-C, pXT7-GFP-Axin2L (originally referred to as XARP) (Alexandrova and Sokol, 2010; Itoh et al., 2000), pCS2-myr-GFP, pCS2-myr-TagBFP-HA (Matsuda et al., 2023), pCS105-HA-RFP-ADIP and pCS2-FlagTurboID-ADIP (Chu et al., 2025). pXT7-GFP-Axin2L was generated from previously described pXT7-GFP-XARP (Alexandrova and Sokol, 2010) by inserting the N-terminal Axin2L sequence encoding aa1-39. This sequence was synthesized as a gBlock fragment by Integrated DNA Technologies and flanked by BspEI and NcoI restriction sites for cloning.

The deletion constructs pXT7-GFP-Axin2L-ΔRGS, pXT7-GFP-Axin2L-ΔGSK3β-binding domain (BD), pXT7-GFP-Axin2L-Δβ-Cat-BD and pXT7-GFP-Axin2L-ΔDIX were generated by PCR using specific primers. The previously described pXT7-GFP-XARP construct is referred to in this study as pXT7-GFP-Axin2L-ΔTBD, because it was missing the Tankyrase-binding domain. All constructs were verified by sequencing, primer details are available upon request. Capped mRNAs were synthesized from linearized plasmid templates using the mMessage mMachine SP6 or T7 transcription kits (Invitrogen) and purified with the RNeasy Mini Kit (Qiagen).

The Axin2L morpholino (Axin2L-MO; 5′-TCAGTACTCCAGCAGAACTCATG-3′) has a 100% match to both Axin2L.L and Axin2L.S genes at the Xenbase and control morpholino (CoMO; 5′-GCTTCAGCTAGTGACACATGCAT-3′) were synthesized by GeneTools.

### Xenopus embryos, microinjections, wound healing assay

Wild-type *Xenopus laevis* were purchased from Xenopus 1 (Michigan) and maintained in accordance with the guidelines of the Institutional Animal Care and Use Committee (IACUC) at the Icahn School of Medicine at Mount Sinai. *In vitro* fertilization and embryo culture were carried out as previously described (Ossipova et al., 2014), and embryonic staging was determined according to (Nieuwkoop and Faber, 1994). Embryos were cultured in 0.1 x Marc’s Modified Ringer’s solution (MMR, 50 mM NaCl, 1 mM KCl, 1 mM CaCl_2_, 0.5 mM MgCl_2_, 2.5 mM HEPES pH 7.4) (Peng, 1991). For microinjections, 2 to 4 cell embryos were transferred to 3% Ficoll in 0.5 x MMR. RNA or morpholino oligonucleotides (MOs) were diluted in RNase-free water (Invitrogen) and 5–10 nl were injected into animal or equatorial sites of four to eight-cell embryos. Amounts of injected mRNA or MO are indicated in figure legends. Injected embryos were transferred to 0.1 x MMR at blastula stages.

Wound healing assays were performed as described previously (Chu et al., 2025). For wound polarity assays, 1.5 ng RNA encoding GFP-tagged Axin2L constructs was injected with a lineage tracer into ventral blastomeres of four-cell embryos. At stage 12, small wounds were generated manually in the embryonic ectoderm using fine dissection forceps. Embryos were allowed to heal for 30-45 min after wounding and were then fixed in 1× MEMFA for imaging.

To validate Axin2L-MO efficiency, four-cell embryos were injected with 20 ng Axin2L-MO into all four blastomeres at the marginal zone. Embryos were collected at stages 12 and 12.5, and lysates were analyzed by immunoblotting with anti-Axin2L antibody. Anti-Dvl2 and anti-α-Actin antibodies were used as specificity and loading controls, respectively.

### Proximity biotinylation, immunoprecipitation and immunoblotting

For proximity biotinylation assays, all four animal blastomeres of four-cell embryos were injected with 200 pg Flag-TurboID-ADIP RNA *(26),* 400 pg GFP-Axin2L RNA and 0.8 mM Biotin (Sigma-Aldrich) as described (Reis et al., 2021). Embryos were cultured until stage 10.5 and lysed for immunoprecipitation using GFP trap (Proteintech). Biotinylated proteins were detected in immunoblotting with anti-Biotin antibody conjugated with HRP (1:1000, Cell Signaling).

Immunoprecipitations and immunoblotting have been carried out as previously described (Itoh et al., 2021). Four-cell embryos were injected animally into each blastomere with RNAs encoding Myc-Axin2L, MycΔRGS-Axin2L (1 ng each) or GFP-Axin2L deletion constructs (1.5 ng). Embryos were lysed in 1 % Triton X-100 lysis buffer (LB) containing 50 mM NaCl, 1 mM EDTA and 50 mM Tris-HCl (pH 7.5). Immunoblotting was carried out with mouse monoclonal anti-Myc (1:200, 9E10 hybridoma supernatant), anti-GFP (1:1000, B2, Santa Cruz), anti-αActin (1:1000, Developmental Studies Hybridoma Bank), anti-Dvl2 (1:500; (Itoh et al., 2005)), or rabbit anti-Axin2L/XARP (1:500, Ab2, (Alexandrova and Sokol, 2010)) antibodies. Chemiluminescence signals were detected using ChemiDoc (BioRad).

### Protein localization, immunostaining and imaging, quantification of cilia length

To assess Axin2L subcellular distribution in the neural or non-neural ectoderm, 400 pg of RNA encoding different GFP-Axin2L constructs and 50 pg of myrBFP mRNA were injected into one dorsal-animal or ventral animal blastomere at the four-cell stage. To examine Axin2L and ADIP colocalization, 300 pg GFP-Axin2L mRNA, 300 pg HA-RFP-ADIP mRNA, and 50 pg myrBFP mRNA were co-injected dorsally at the four-cell stage. For F-actin visualization, embryos were fixed in MEMFA for 30 min at room temperature. Fixed embryos were permeabilized with 0.1% Triton X-100 in PBS for 10 min and incubated overnight at 4°C with Alexa Fluor 555-conjugated phalloidin (1:500; Thermo Fisher Scientific) diluted in PBS containing 1% BSA.

For Vangl2 polarization assays, four-cell embryos were injected into one dorsal animal blastomere with 500 pg RNA encoding myc-Axin2L or myc-Axin2L-ΔRGS and HA-Axin2L-C. For knockdown and rescue experiments, one dorsal-animal blastomere was injected with either 30 ng Axin2L-MO or 30 ng CoMO together with 10 pg HA-myrBFP RNA as a lineage tracer. For rescue, 30 ng Axin2L-MO was coinjected with 500 pg Axin2L-ΔTBD RNA and 10 pg HA-myrBFP RNA as a lineage tracer. Injected embryos were cultured in 0.1× MMR at 12-14°C until the desired developmental stage. Each experimental condition included at least 15–20 embryos, and all experiments were repeated independently at least three times.

For Myc, HA, and Vangl2 immunostaining, stage 13 embryos were devitellinized in 0.1× MMR and fixed in cold 2% trichloroacetic acid (TCA) for 15 min at room temperature. Embryos were washed three times in 1× PBS with 0.1% TritonX100 (PBST) for 5 min each and blocked for 1 h at room temperature in 20% fetal bovine serum (FBS) in 1× PBST. Embryos were then incubated overnight at 4°C with primary antibodies diluted in 10% FBS in 1× PBST. Primary antibodies included rabbit anti-Vangl2 antibody (1:200 (Ossipova et al., 2015)) and mouse anti-HA antibody (12CA5, 1:200) to detect myrBFP-HA lineage tracer in MO-injected cells and mouse anti-Myc antibody (9E10, 1:200) to detect myc-Axin2L constructs. After primary antibody incubation, embryos were washed five times in 1× PBST at room temperature, with 1 h intervals between washes. Embryos were then incubated overnight at 4°C with secondary antibodies diluted in 10% FBS in 1× PBST. Secondary antibodies included Alexa Fluor 488-conjugated goat anti-mouse IgG (1:300; Thermo Fisher Scientific) and Cy3-conjugated donkey anti-rabbit IgG (1:400; Jackson ImmunoResearch). Immunostained embryos were washed three times in PBST for 10 min each before imaging. Although Vangl2 and Myc/HA signals were acquired according to the fluorophores listed above, image channels were pseudocolored in the figures for clarity and color-blind-friendly presentation.

For gastrocoel roof plate (GRP) cilia analysis, four-cell embryos were injected with 40 ng Axin2L MO and 10 pg myrGFP RNA into one dorsal blastomere at the medial equator. When sibling embryos reached stage 17, the injected embryos were fixed in 3.7% formaldehyde/PBS. GRP explants were dissected and stained with rabbit anti-GFP (1:300, A6455, Invitrogen) and mouse anti-acetylated-tubulin (1:1000, 6-11B1, sc-23950, Santa Cruz Biotechnology) primary antibodies and Alexa488-conjugated anti-rabbit IgG (1:250, Invitrogen) and Cy3-conjugated anti-mouse IgG (Jackson ImmunoResearch, 1:200) secondary antibodies.

Fluorescence imaging was performed using BC43 spinning disk confocal microscope (Andor, Oxford Instruments) with a 20× objective lens and the Fusion software version 2. Z-stacks were acquired and processed as maximum-intensity projections using ImageJ for subsequent analysis and quantification. Each experiment was repeated at least three times, with a minimum of ten embryos per experimental group unless otherwise indicated. Phenotypes and localization patterns described in the text were reproducible across independent experiments.

To quantify GRP cilia length, the stained cilia were manually outlined using a free-hand line tool in ImageJ software. Length of cilia in control GRP (no GFP) was compared with cilia in the GRP of GFP-positive Axin2L-depleted cells. Student’s t-test was used to determine statistical significance between the two groups. ****p<0.0001.

### Polarity quantification and statistical analysis

Axial defects of embryos were quantified and statistically analyzed using GraphPad Prism v11.0.2. Axin2L and ADIP polarity were quantified as described (Santhi Velayudhan et al., 2026), using a Python-based workflow available at https://github.com/sujaynelson/Endogenous-ADIP-PCP-Quantification.git. Protein localization was quantified using either puncta-based or diffuse pixel-based analysis, depending on the observed distribution of each Axin2L construct. For punctate proteins, fluorescence images were log-transformed, normalized, and thresholded to identify clusters. Cell polarity vectors were calculated from the cell centroid toward the cluster centroids, weighted by cluster size or fluorescence intensity. For diffusely localized Axin2L-ΔDIX, membrane-adjacent pixels were excluded, and background-subtracted cytoplasmic fluorescence was used to calculate an intensity-weighted centroid. A polarity vector was then drawn from the cell centroid to this intensity-weighted centroid. A uniform-cytoplasm control was used to estimate potential bias caused by cell shape or segmentation. In both analyses, vector orientations were compared with the relevant reference direction, such as the wound-facing axis or neural plate border-facing axis, displayed as rose plots, and evaluated using Chi-square tests of angular distributions.

Vangl2 polarity was quantified using Python scripts implementing a quadrant-ratio analysis adapted from (Hirano et al., 2022), using the Python workflow available at https://github.com/sierhill/Vangl2-Polarity-Quantification-Quadrant-Ratio. Cellpose segmentation masks (Stringer et al., 2021) were used to define individual cell boundaries, and Vangl2 fluorescence intensity was measured along the one-pixel-wide membrane boundary of each cell. Vangl2 fluorescence intensity measured along the cell boundary was converted into a 360-bin angular intensity profile centered on the cell centroid. For each cell, all possible axes from 0° to 179° were tested by summing Vangl2 intensity in two opposite 90° quadrants and comparing this value with the summed intensity in the perpendicular quadrants. The polarity value was defined as the ratio between opposite-quadrant pairs. Because opposite quadrants were summed, polarity was analyzed as an undirected axis. For two-channel datasets, Vangl2 polarity was quantified from the Vangl2 channel, whereas membrane or lineage markers, when present, quantified in a separate channel. Polarity-axis distributions are displayed as angular rose plots. Mean polarity values between groups are shown as AP-versus-ML ratios and compared using two-sided Mann–Whitney test.

### Axin2L-ADIP puncta colocalization analysis

Axin2L-ADIP puncta colocalization was quantified using the ComDet plugin in Fiji (Schindelin et al., 2012). ADIP was assigned to channel 1 and Axin2L to channel 2. Puncta were detected independently in each channel using the particle size of 2 pixels and an intensity threshold of 4-6 standard deviations above the local background. Larger-particle inclusion and segmentation options were enabled. Puncta with centers located within 8 pixels of each other were classified as colocalized. Each matched ADIP-Axin2L puncta pair was counted once. Unmatched detections were classified as non-colocalized Axin2L puncta or non-colocalized ADIP puncta, respectively. Background-corrected integrated puncta intensities from both channels were used for scatter plot visualization in Microsoft Excel.

## Acknowledgments

We thank Sujay Nelson for help with quantification of Vangl2 planar polarity. We also thank Rodrigo Fernandez-Gonzalez for comments on the manuscript. We are grateful to members of the Sokol laboratory for valuable discussions. This research was supported by the NIH grants R35GM122492 and R01HD092990 to S.Y.S.

## Author contributions

S.S.V. designed, executed, and analyzed most of the experiments, prepared figures, and wrote the manuscript. K.I. showed the physical interaction of Axin2L and ADIP, characterized embryonic phenotypes and cilia formation, validated Axin2L antibodies and morpholinos. S.Y.S. conceived the study, designed experiments, acquired funding, supervised the project and wrote the manuscript. All authors agreed on the final version of the manuscript.

## Data availability

Raw data and images have been deposited in the Mendeley Data.

## Supplementary material

**Fig. S1.**
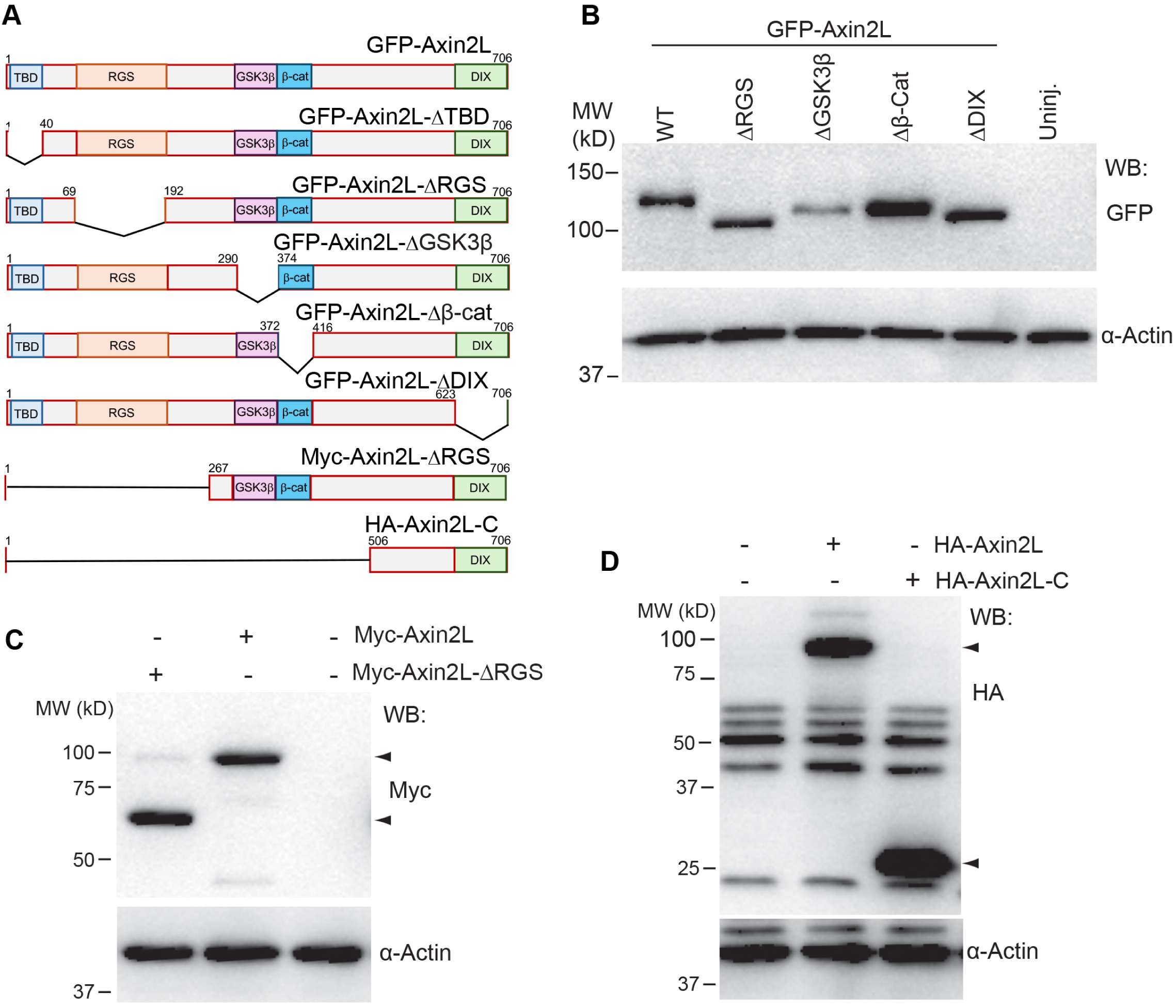
Axin2L constructs and their expression levels in *Xenopus* gastrulae. (**A**) Scheme of GFP-Axin2L constructs used in this study. GFP-Axin2L contains the Tankyrase-binding domain (TBD), the RGS domain, the GSK3β-binding and β-catenin-binding regions, and the DIX domain. (**B-D**) Four to eight cell embryos were injected 4 times animally with RNA encoding GFP-Axin2L constructs (1.5 ng), Myc-Axin2L, Myc-Axin2L-ΔRGS (1 ng), HA-Axin2L and HA-Axin2L-C (2 ng). α−Actin serves as loading control. (**B**) Immunoblotting of lysates from embryos expressing different GFP-Axin2L deletion constructs, using anti-GFP antibody. **C, D**, Comparable expression of Myc-Axin2L and Myc-Axin2L-ΔRGS (**C**), and HA-Axin2L and HA-Axin2L-C (**D**) proteins in embryo lysates.

**Fig. S2.**
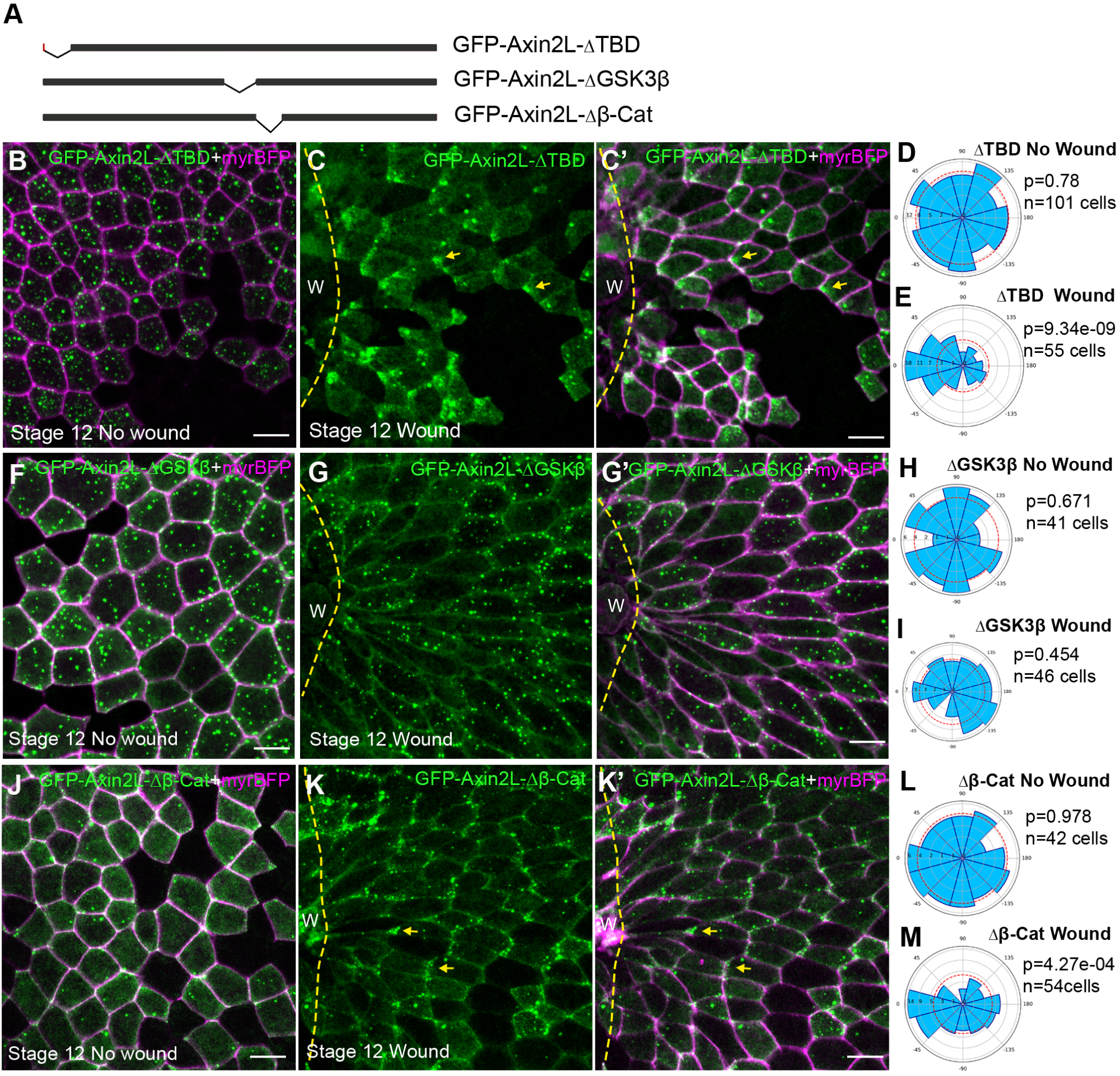
The GSK3-binding domain is required for Axin2L polarization during wound healing. (**A**) Scheme of Axin2L constructs used in this experiment. GFP-Axin2L-ΔTBD lacks the Tankyrase-binding domain, GFP-Axin2L-ΔGSK3β lacks the GSK3β-binding region, and GFP-Axin2L-Δβ-Cat lacks the β-catenin–binding region. Four-cell stage embryos were injected with 1.5 ng of RNA encoding GFP-Axin2L-ΔTBD, GFP-Axin2L-ΔGSK3β-binding domain, or GFP-Axin2L-Δβ-Cat-BD together with 50 pg of myrBFP RNA as a cell border tracer. Stage 12 (st.12) surface ectoderm was imaged intact or at 45 min after wounding. GFP-Axin2L constructs are in green, and myrBFP-labeled cell borders are in magenta. Yellow dashed lines indicate wound edge. (**B**) GFP-Axin2L-ΔTBD puncta in control unwounded ectoderm. (**C, C′**) After wounding, GFP-Axin2L-ΔTBD becomes polarized toward wound edge. (**D, E**) Rose plots quantify data shown in B, C. (**F, G**) Lack of polarization of GFP-Axin2L-ΔGSK3β puncta in both control (**F**) and wounded (**G, G**’) ectoderm. (**H, I**) Rose plots quantify data in (F, G). (**J**) GFP-Axin2L-Δβ-Cat puncta do not polarize in control ectoderm. (K, K′) After wounding, GFP-Axin2L-Δβ-Cat puncta polarize toward wound edge. (**L, M**) Rose plots quantify data shown in J, K. In (**D, E, H, I, L, M**), rose plots quantify Axin2L puncta orientation relative to wound edge or the reference axis in unwounded controls. Chi-square test, data are from three embryos per group, three experiments.

**Fig. S3.**
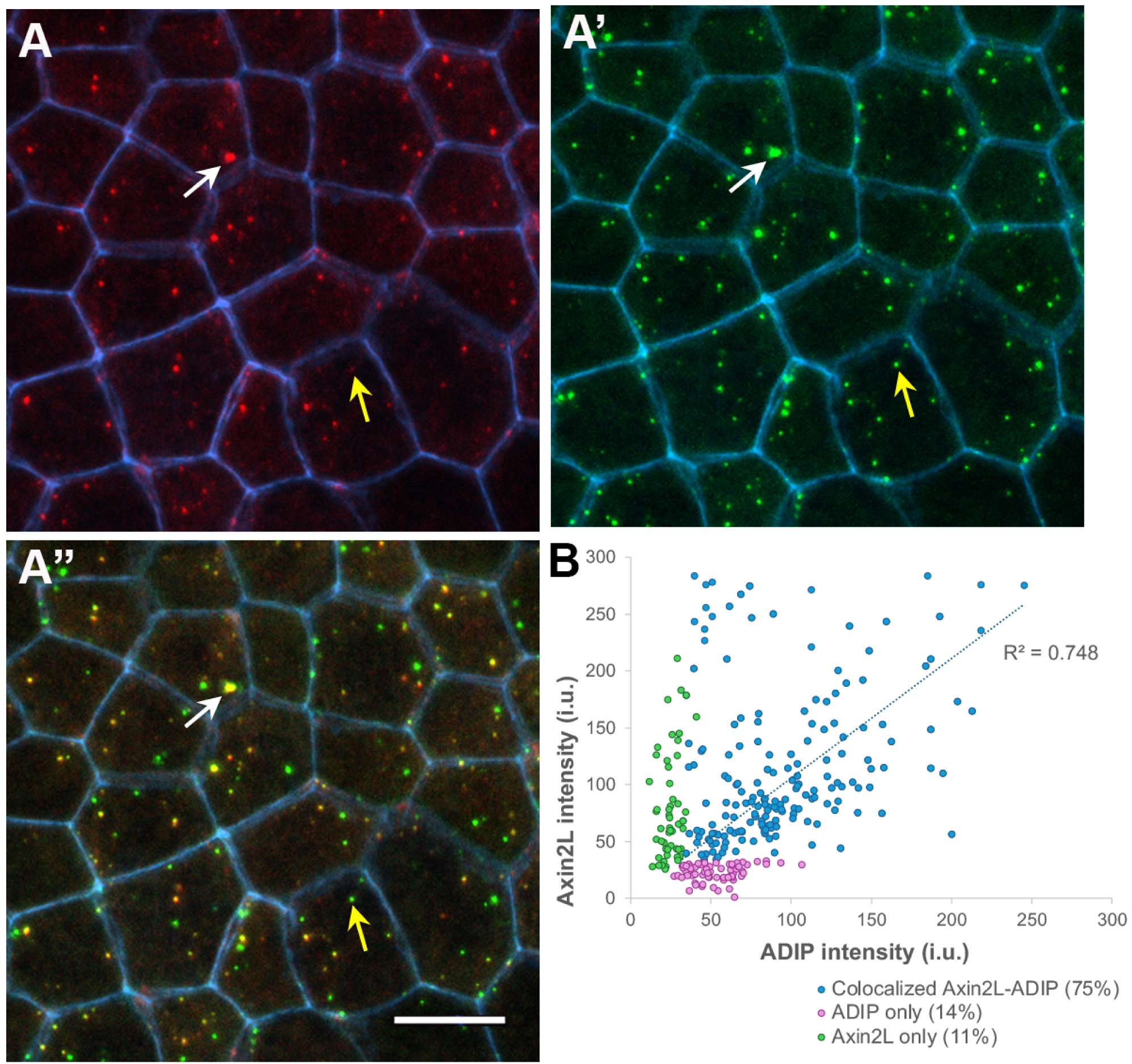
Colocalization analysis of ADIP and Axin2L puncta. (**A-A″**) Representative stage 12 ectodermal cells coexpressing HA-RFP-ADIP (300 pg, red), GFP-Axin2L (300 pg, green), and myrBFP membrane tracer (50 pg, blue). (**A**) HA-RFP-ADIP puncta. (**A′**) GFP-Axin2L puncta. (**A″**) Merged image showing overlap between ADIP and Axin2L puncta. Yellow arrows indicate colocalized ADIP and Axin2L puncta with partial overlap or unequal relative intensities, whereas white arrows mark strongly overlapping puncta at cell junctions. Scale bar, 20 μm. (**B**) Scatter plot showing integrated puncta intensity for ADIP and Axin2L. Colocalized ADIP-Axin2L puncta (75%) are shown in blue, non-colocalized Axin2L puncta in magenta, and non-colocalized ADIP puncta in green. Linear regression is indicated by the dashed line. N = 2 embryos.

**Fig. S4.**
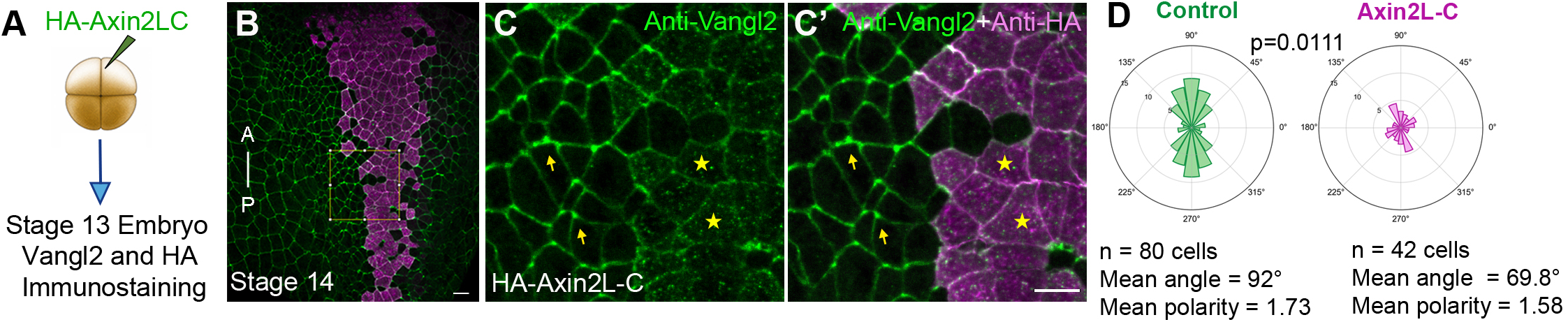
Axin2L-C interferes with planar cell polarity in the neuroectoderm. (**A**) Four-cell embryos were unilaterally injected with 500 pg HA-Axin2L-C RNA in the dorsal region, fixed at stage 13 and immunostained for Vangl2 and HA. (**B**) Representative stage 13 neuroectoderm mosaically expressing HA-Axin2L-C. Anteroposterior (A–P) axis is indicated. (**C, C′**) Higher magnification of the boxed region in (**B**) shows Vangl2 staining (green) in control uninjected cells and HA-Axin2L-C-expressing cells (**C**), marked by anti-HA staining in **C’** (magenta). Arrows point to polarized Vangl2 in uninjected cells, whereas asterisks mark reduced or absent Vangl2 polarization. (B′) Merged image. Scale bars, 20 μm. (**D**) Quantification of cells displaying AP-vs-ML axis Vangl2 orientation. Axin2L-C expression significantly reduces Vangl2 polarization compared with control cells (Mann-Whitney, p = 0.0111). Data from three embryos across three independent experiments.

**Fig. S5.**
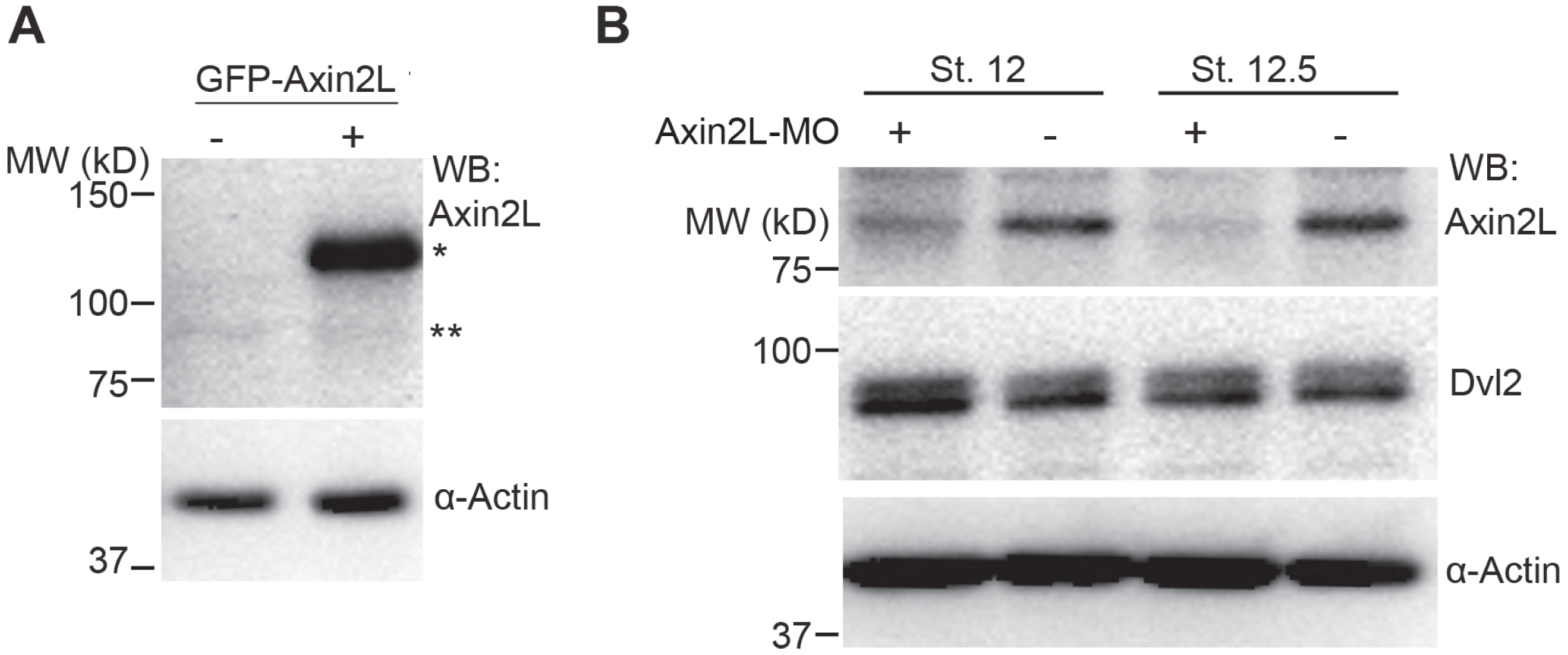
Validation of Axin2L antibody specificity and confirmation of endogenous Axin2L knockdown. (**A**) Four-cell embryos were injected with GFP-Axin2lΔTBD RNA (400 pg) into each blastomere. Lysates were prepared from injected stage 11 embryos and uninjected siblings for immunoblotting with anti-Axin2L and anti-α-Actin antibodies to control for loading. Overexpressed GFP-Axin2L band is indicated with *, while endogenous Axin2L is marked with **. (**B**) Embryos were injected with 20 ng Axin2L MO into all four blastomeres at the marginal zone. Immunoblotting was performed with lysates collected when control embryos reached stages 12 and 12.5, using anti-Axin2L, anti-Dvl2, and anti-αActin to control for loading. Specific reduction of the 85 kDa protein band of Axin2L is detected in MO-injected embryo lysates. Three experiments with 5-10 embryos per group.

## REFERENCES

Aceves-Ewing, N.M., Lanza, D.G., Marcogliese, P.C., Lu, D., Hsu, C.W., Gonzalez, M., Christiansen, A.E., Rasmussen, T.L., Ho, A.J., Gaspero, A., Seavitt, J., Dickinson, M.E., Yuan, B., Shayota, B.J., Pachter, S., Hu, X., Day-Salvatore, D.L., Mackay, L., Kanca, O., Wangler, M.F., Potocki, L., Rosenfeld, J.A., Lewis, R.A., Chao, H.T., Lee, B., Lee, S., Undiagnosed Diseases, N., Baylor College of Medicine Center for Precision Medicine, M., Yamamoto, S., Bellen, H.J., Burrage, L.C., Heaney, J.D., 2025. Uncovering Phenotypic Expansion in AXIN2-Related Disorders through Precision Animal Modeling. medRxiv.

Adler, P.N., Wallingford, J.B., 2017. From Planar Cell Polarity to Ciliogenesis and Back: The Curious Tale of the PPE and CPLANE proteins. Trends Cell Biol 27, 379–390.

Alexandrova, E.M., Sokol, S.Y., 2010. Xenopus axin-related protein: a link between its centrosomal localization and function in the Wnt/beta-catenin pathway. Dev Dyn 239, 261–270.

Asada, M., Irie, K., Morimoto, K., Yamada, A., Ikeda, W., Takeuchi, M., Takai, Y., 2003. ADIP, a novel Afadin- and alpha-actinin-binding protein localized at cell-cell adherens junctions. J Biol Chem 278, 4103–4111.

Azzolin, L., Panciera, T., Soligo, S., Enzo, E., Bicciato, S., Dupont, S., Bresolin, S., Frasson, C., Basso, G., Guzzardo, V., Fassina, A., Cordenonsi, M., Piccolo, S., 2014. YAP/TAZ Incorporation in the beta-Catenin Destruction Complex Orchestrates the Wnt Response. Cell 158, 157–170.

Behrens, J., Jerchow, B.A., Wurtele, M., Grimm, J., Asbrand, C., Wirtz, R., Kuhl, M., Wedlich, D., Birchmeier, W., 1998. Functional interaction of an axin homolog, conductin, with beta-catenin, APC, and GSK3beta. Science 280, 596–599.

Blum, M., Feistel, K., Thumberger, T., Schweickert, A., 2014. The evolution and conservation of left-right patterning mechanisms. Development 141, 1603–1613.

Branon, T.C., Bosch, J.A., Sanchez, A.D., Udeshi, N.D., Svinkina, T., Carr, S.A., Feldman, J.L., Perrimon, N., Ting, A.Y., 2018. Efficient proximity labeling in living cells and organisms with TurboID. Nat Biotechnol 36, 880–887.

Butler, M.T., Wallingford, J.B., 2017. Planar cell polarity in development and disease. Nat Rev Mol Cell Biol 18, 375–388.

Caddy, J., Wilanowski, T., Darido, C., Dworkin, S., Ting, S.B., Zhao, Q., Rank, G., Auden, A., Srivastava, S., Papenfuss, T.A., Murdoch, J.N., Humbert, P.O., Parekh, V., Boulos, N., Weber, T., Zuo, J., Cunningham, J.M., Jane, S.M., 2010. Epidermal wound repair is regulated by the planar cell polarity signaling pathway. Dev Cell 19, 138–147.

Castellone, M.D., Teramoto, H., Williams, B.O., Druey, K.M., Gutkind, J.S., 2005. Prostaglandin E2 promotes colon cancer cell growth through a Gs-axin-beta-catenin signaling axis. Science 310, 1504–1510.

Cervenka, I., Valnohova, J., Bernatik, O., Harnos, J., Radsetoulal, M., Sedova, K., Hanakova, K., Potesil, D., Sedlackova, M., Salasova, A., Steinhart, Z., Angers, S., Schulte, G., Hampl, A., Zdrahal, Z., Bryja, V., 2016. Dishevelled is a NEK2 kinase substrate controlling dynamics of centrosomal linker proteins. Proc Natl Acad Sci U S A 113, 9304–9309.

Chu, C.W., Velayudhan, S.S., Schauser, J.H., Krishnakumar, S., Yang, S., Itoh, K., Alfandari, D., Trusina, A., Sokol, S.Y., 2025. Mechanical cues polarize ADIP protein complexes to control vertebrate morphogenesis and wound healing. Curr. Biol. 35, 3315–3326 e3314.

Devenport, D., 2014. The cell biology of planar cell polarity. The Journal of cell biology 207, 171–179.

Fagotto, F., Jho, E., Zeng, L., Kurth, T., Joos, T., Kaufmann, C., Costantini, F., 1999. Domains of axin involved in protein-protein interactions, Wnt pathway inhibition, and intracellular localization. J Cell Biol 145, 741–756.

Fumoto, K., Kadono, M., Izumi, N., Kikuchi, A., 2009. Axin localizes to the centrosome and is involved in microtubule nucleation. EMBO Rep 10, 606–613.

Goodrich, L.V., Strutt, D., 2011. Principles of planar polarity in animal development. Development 138, 1877–1892.

Gray, R.S., Roszko, I., Solnica-Krezel, L., 2011. Planar cell polarity: coordinating morphogenetic cell behaviors with embryonic polarity. Dev Cell 21, 120–133.

Hart, M.J., de los Santos, R., Albert, I.N., Rubinfeld, B., Polakis, P., 1998. Downregulation of beta-catenin by human Axin and its association with the APC tumor suppressor, beta-catenin and GSK3 beta. Curr Biol 8, 573–581.

Hashimoto, M., Shinohara, K., Wang, J., Ikeuchi, S., Yoshiba, S., Meno, C., Nonaka, S., Takada, S., Hatta, K., Wynshaw-Boris, A., Hamada, H., 2010. Planar polarization of node cells determines the rotational axis of node cilia. Nat Cell Biol 12, 170–176.

Hikasa, H., Sokol, S.Y., 2013. Wnt signaling in vertebrate axis specification. Cold Spring Harb Perspect Biol 5, a007955.

Hirano, S., Mii, Y., Charras, G., Michiue, T., 2022. Alignment of the cell long axis by unidirectional tension acts cooperatively with Wnt signalling to establish planar cell polarity. Development 149.

Hori, A., Ikebe, C., Tada, M., Toda, T., 2014. Msd1/SSX2IP-dependent microtubule anchorage ensures spindle orientation and primary cilia formation. EMBO Rep 15, 175–184.

Hori, A., Peddie, C.J., Collinson, L.M., Toda, T., 2015. Centriolar satellite- and hMsd1/SSX2IP-dependent microtubule anchoring is critical for centriole assembly. Mol Biol Cell 26, 2005–2019.

Ikeda, S., Kishida, S., Yamamoto, H., Murai, H., Koyama, S., Kikuchi, A., 1998. Axin, a negative regulator of the Wnt signaling pathway, forms a complex with GSK-3beta and beta-catenin and promotes GSK-3beta-dependent phosphorylation of beta-catenin. EMBO J 17, 1371–1384.

Itoh, K., Antipova, A., Ratcliffe, M.J., Sokol, S., 2000. Interaction of dishevelled and Xenopus axin-related protein is required for wnt signal transduction. Mol Cell Biol 20, 2228–2238.

Itoh, K., Brott, B.K., Bae, G.U., Ratcliffe, M.J., Sokol, S.Y., 2005. Nuclear localization is required for Dishevelled function in Wnt/beta-catenin signaling. J Biol 4, 3.

Itoh, K., Jenny, A., Mlodzik, M., Sokol, S.Y., 2009. Centrosomal localization of Diversin and its relevance to Wnt signaling. J Cell Sci 122, 3791–3798.

Itoh, K., Krupnik, V.E., Sokol, S.Y., 1998. Axis determination in Xenopus involves biochemical interactions of axin, glycogen synthase kinase 3 and beta-catenin. Curr Biol 8, 591–594.

Itoh, K., Ossipova, O., Sokol, S.Y., 2021. Pinhead antagonizes Admp to promote notochord formation. iScience 24, 102520.

Kang, K., Shi, Q., Wang, X., Chen, Y.G., 2022. Dishevelled phase separation promotes Wnt signalosome assembly and destruction complex disassembly. J Cell Biol 221.

Kishida, S., Yamamoto, H., Hino, S., Ikeda, S., Kishida, M., Kikuchi, A., 1999. DIX domains of Dvl and axin are necessary for protein interactions and their ability to regulate beta-catenin stability. Mol Cell Biol 19, 4414–4422.

Komiya, Y., Habas, R., 2008. Wnt signal transduction pathways. Organogenesis 4, 68–75.

Lach, R.S., Qiu, C., Kajbaf, E.Z., Baxter, N., Han, D., Wang, A., Lock, H., Chirikian, O., Pruitt, B., Wilson, M.Z., 2022. Nucleation of the destruction complex on the centrosome accelerates degradation of beta-catenin and regulates Wnt signal transmission. Proc Natl Acad Sci U S A 119, e2204688119.

MacDonald, B.T., Tamai, K., He, X., 2009. Wnt/beta-catenin signaling: components, mechanisms, and diseases. Dev Cell 17, 9–26.

Matsuda, M., Rozman, J., Ostvar, S., Kasza, K.E., Sokol, S.Y., 2023. Mechanical control of neural plate folding by apical domain alteration. Nat Commun 14, 8475.

Matsuda, M., Sokol, S.Y., 2021. Xenopus neural tube closure: A vertebrate model linking planar cell polarity to actomyosin contractions. Curr Top Dev Biol 145, 41–60.

Maurice, M.M., Angers, S., 2025. Mechanistic insights into Wnt-beta-catenin pathway activation and signal transduction. Nat Rev Mol Cell Biol 26, 371–388.

McNeill, H., 2010. Planar cell polarity: keeping hairs straight is not so simple. Cold Spring Harb Perspect Biol 2, a003376.

Miete, C., Solis, G.P., Koval, A., Bruckner, M., Katanaev, V.L., Behrens, J., Bernkopf, D.B., 2022. Galphai2-induced conductin/axin2 condensates inhibit Wnt/beta-catenin signaling and suppress cancer growth. Nat Commun 13, 674.

Niehrs, C., Da Silva, F., Seidl, C., 2025. Cilia as Wnt signaling organelles. Trends Cell Biol 35, 24–32.

Nieuwkoop, P.D., Faber, J., 1994. Normal table of Xenopus laevis (Daudin): a systematical and chronological survey of the development from the fertilized egg till the end of metamorphosis. Garland Pub., New York.

Nikolopoulou, E., Galea, G.L., Rolo, A., Greene, N.D., Copp, A.J., 2017. Neural tube closure: cellular, molecular and biomechanical mechanisms. Development 144, 552–566.

Nong, J., Kang, K., Shi, Q., Zhu, X., Tao, Q., Chen, Y.G., 2021. Phase separation of Axin organizes the beta-catenin destruction complex. J Cell Biol 220.

Nusse, R., Clevers, H., 2017. Wnt/beta-Catenin Signaling, Disease, and Emerging Therapeutic Modalities. Cell 169, 985–999.

Ossipova, O., Chuykin, I., Chu, C.W., Sokol, S.Y., 2015. Vangl2 cooperates with Rab11 and Myosin V to regulate apical constriction during vertebrate gastrulation. Development 142, 99–107.

Ossipova, O., Kim, K., Lake, B.B., Itoh, K., Ioannou, A., Sokol, S.Y., 2014. Role of Rab11 in planar cell polarity and apical constriction during vertebrate neural tube closure. Nat Commun 5, 3734.

Park, T.J., Haigo, S.L., Wallingford, J.B., 2006. Ciliogenesis defects in embryos lacking inturned or fuzzy function are associated with failure of planar cell polarity and Hedgehog signaling. Nat Genet 38, 303–311.

Park, T.J., Mitchell, B.J., Abitua, P.B., Kintner, C., Wallingford, J.B., 2008. Dishevelled controls apical docking and planar polarization of basal bodies in ciliated epithelial cells. Nat Genet 40, 871–879.

Peng, H.B., 1991. Xenopus laevis: Practical uses in cell and molecular biology. Solutions and protocols. Methods Cell Biol 36, 657–662.

Peng, Y., Axelrod, J.D., 2012. Asymmetric protein localization in planar cell polarity: mechanisms, puzzles, and challenges. Curr Top Dev Biol 101, 33–53.

Qian, L., Mahaffey, J.P., Alcorn, H.L., Anderson, K.V., 2011. Tissue-specific roles of Axin2 in the inhibition and activation of Wnt signaling in the mouse embryo. Proc Natl Acad Sci U S A 108, 8692–8697.

Reis, A.H., Xiang, B., Ossipova, O., Itoh, K., Sokol, S.Y., 2021. Identification of the centrosomal maturation factor SSX2IP as a Wtip-binding partner by targeted proximity biotinylation. PLoS One 16, e0259068.

Santhi Velayudhan, S., Itoh, K., Chu, C.W., Alfandari, D., Sokol, S.Y., 2026. Planar polarization of endogenous ADIP during Xenopus neurulation. Biol Open 15.

Schaefer, K.N., Bonello, T.T., Zhang, S., Williams, C.E., Roberts, D.M., McKay, D.J., Peifer, M., 2018. Supramolecular assembly of the beta-catenin destruction complex and the effect of Wnt signaling on its localization, molecular size, and activity in vivo. PLoS Genet 14, e1007339.

Schindelin, J., Arganda-Carreras, I., Frise, E., Kaynig, V., Longair, M., Pietzsch, T., Preibisch, S., Rueden, C., Saalfeld, S., Schmid, B., Tinevez, J.Y., White, D.J., Hartenstein, V., Eliceiri, K., Tomancak, P., Cardona, A., 2012. Fiji: an open-source platform for biological-image analysis. Nat Methods 9, 676–682.

Schlessinger, K., McManus, E.J., Hall, A., 2007. Cdc42 and noncanonical Wnt signal transduction pathways cooperate to promote cell polarity. J Cell Biol 178, 355–361.

Schubert, A., Voloshanenko, O., Ragaller, F., Gmach, P., Kranz, D., Scheeder, C., Miersch, T., Schulz, M., Trumper, L., Binder, C., Lampe, M., Engel, U., Boutros, M., 2022. Superresolution microscopy localizes endogenous Dvl2 to Wnt signaling-responsive biomolecular condensates. Proc Natl Acad Sci U S A 119, e2122476119.

Schwarz-Romond, T., Asbrand, C., Bakkers, J., Kuhl, M., Schaeffer, H.J., Huelsken, J., Behrens, J., Hammerschmidt, M., Birchmeier, W., 2002. The ankyrin repeat protein Diversin recruits Casein kinase Iepsilon to the beta-catenin degradation complex and acts in both canonical Wnt and Wnt/JNK signaling. Genes Dev 16, 2073–2084.

Semenov, M.V., Habas, R., Macdonald, B.T., He, X., 2007. SnapShot: Noncanonical Wnt Signaling Pathways. Cell 131, 1378.

Shnitsar, I., Bashkurov, M., Masson, G.R., Ogunjimi, A.A., Mosessian, S., Cabeza, E.A., Hirsch, C.L., Trcka, D., Gish, G., Jiao, J., Wu, H., Winklbauer, R., Williams, R.L., Pelletier, L., Wrana, J.L., Barrios-Rodiles, M., 2015. PTEN regulates cilia through Dishevelled. Nat Commun 6, 8388.

Smalley, M.J., Sara, E., Paterson, H., Naylor, S., Cook, D., Jayatilake, H., Fryer, L.G., Hutchinson, L., Fry, M.J., Dale, T.C., 1999. Interaction of axin and Dvl-2 proteins regulates Dvl-2-stimulated TCF-dependent transcription. EMBO J 18, 2823–2835.

Sokol, S.Y., 1996. Analysis of Dishevelled signalling pathways during Xenopus development. Current Biology 6, 1456–1467.

Sokol, S.Y., 2015. Spatial and temporal aspects of Wnt signaling and planar cell polarity during vertebrate embryonic development. Semin Cell Dev Biol 42, 78–85.

Stringer, C., Wang, T., Michaelos, M., Pachitariu, M., 2021. Cellpose: a generalist algorithm for cellular segmentation. Nat Methods 18, 100–106.

Tada, M., Smith, J.C., 2000. Xwnt11 is a target of Xenopus Brachyury: regulation of gastrulation movements via Dishevelled, but not through the canonical Wnt pathway. Development 127, 2227–2238.

Takeshita, H., Sawa, H., 2005. Asymmetric cortical and nuclear localizations of WRM-1/beta-catenin during asymmetric cell division in C. elegans. Genes Dev 19, 1743–1748.

Torban, E., Sokol, S.Y., 2021. Planar cell polarity pathway in kidney development, function and disease. Nat Rev Nephrol.

Vamadevan, V., Chaudhary, N., Maddika, S., 2022. Ubiquitin-assisted phase separation of dishevelled-2 promotes Wnt signalling. J Cell Sci 135.

van Dop, M., Fiedler, M., Mutte, S., de Keijzer, J., Olijslager, L., Albrecht, C., Liao, C.Y., Janson, M.E., Bienz, M., Weijers, D., 2020. DIX Domain Polymerization Drives Assembly of Plant Cell Polarity Complexes. Cell 180, 427–439 e412.

Velayudhan, S.S., Chu, C.W., Itoh, K., Sokol, S.Y., 2025. Mechanosensitive localization of Diversin highlights its function in vertebrate morphogenesis and planar cell polarity. Biol Open 14.

Wallingford, J.B., Mitchell, B., 2011. Strange as it may seem: the many links between Wnt signaling, planar cell polarity, and cilia. Genes Dev 25, 201–213.

Wallingford, J.B., Rowning, B.A., Vogeli, K.M., Rothbacher, U., Fraser, S.E., Harland, R.M., 2000. Dishevelled controls cell polarity during Xenopus gastrulation. Nature 405, 81–85.

Weber, U., Farhadifar, R., Mlodzik, M., 2025. The Wnt co-receptor Arrow-LRP5/6 is required for Planar Cell Polarity establishment in Drosophila. bioRxiv.

Weiner, A.T., Seebold, D.Y., Torres-Gutierrez, P., Folker, C., Swope, R.D., Kothe, G.O., Stoltz, J.G., Zalenski, M.K., Kozlowski, C., Barbera, D.J., Patel, M.A., Thyagarajan, P., Shorey, M., Nye, D.M.R., Keegan, M., Behari, K., Song, S., Axelrod, J.D., Rolls, M.M., 2020. Endosomal Wnt signaling proteins control microtubule nucleation in dendrites. PLoS Biol 18, e3000647.

Williams, M.L.K., Solnica-Krezel, L., 2020. Cellular and molecular mechanisms of convergence and extension in zebrafish. Curr Top Dev Biol 136, 377–407.

Yasunaga, T., Itoh, K., Sokol, S.Y., 2011. Regulation of basal body and ciliary functions by Diversin. Mech Dev 128, 376–386.

Yu, H.M., Jerchow, B., Sheu, T.J., Liu, B., Costantini, F., Puzas, J.E., Birchmeier, W., Hsu, W., 2005. The role of Axin2 in calvarial morphogenesis and craniosynostosis. Development 132, 1995–2005.

Zallen, J.A., 2007. Planar polarity and tissue morphogenesis. Cell 129, 1051–1063.

Zeng, L., Fagotto, F., Zhang, T., Hsu, W., Vasicek, T.J., Perry, W.L., 3rd, Lee, J.J., Tilghman, S.M., Gumbiner, B.M., Costantini, F., 1997. The mouse Fused locus encodes Axin, an inhibitor of the Wnt signaling pathway that regulates embryonic axis formation. Cell 90, 181–192.

